# GCN5 Maintains Muscle Integrity by Acetylating YY1 to Promote Dystrophin Expression

**DOI:** 10.1101/2021.03.29.436986

**Authors:** Gregory C. Addicks, Hongbo Zhang, Dongryeol Ryu, Goutham Vasam, Philip L. Marshall, Alex E. Green, Sonia Patel, Baeki E. Kang, Doyoun Kim, Elena Katsyuba, Evan G. Williams, Jean-Marc Renaud, Johan Auwerx, Keir J. Menzies

## Abstract

This work identifies a novel role for the acetyltransferase GCN5 in regulating muscle integrity through inhibition of DNA binding activity of the transcriptional repressor YY1. Here we report that in mice a muscle-specific knockout of GCN5 (*Gcn5*^skm-/-^) reduces the expression of key structural muscle proteins, including dystrophin, resulting in myopathy. Supporting our observation, a meta-analysis between the differential transcriptome of *Gcn5*^skm-/-^ muscle and all available open-access data sets identified top correlations with musculoskeletal diseases in humans. GCN5 was found to acetylate YY1 at two residues (K392 and K393), which disrupts the interaction between the YY1 zinc-finger region and DNA. De/acetylation mimics for these YY1 post-translational modifications modulated muscle structural gene expression and DNA binding. Analysis of human GTEx data also found positive and negative correlations between fiber diameter and GCN5 and YY1 respectively. Collectively, our results demonstrate that GCN5 acetyltransferase activity regulates YY1 DNA binding and expression of dystrophin to modulate muscle integrity.

## Introduction

Skeletal muscle is a dynamic tissue that can coordinate structural, metabolic, and contractile adaptations to demands for contractile work. Profound remodeling of muscle can be evoked by physiological signals, including resistance or endurance training, or pathological signals that arise during disease, myopathy or aging. Understanding the molecular signalling pathways that regulate muscle are therefore integral for our understanding of motility throughout the lifespan. Essential for muscle integrity, a critical structural link between the actin cytoskeleton and the extracellular matrix is maintained by the dystrophin-associated protein complex (DAPC), which stabilizes the sarcolemma during contractions, and transmits force generated by the contractile apparatus to the extracellular matrix (Petrof et al., 1993). The DAPC includes intracellular (dystrophin, syncoilin, syntrophin, α-dystrobrevin, and nNOS), transmembrane (β-dystroglycan, sarcoglycans, and sarcospan) and extracellular (α-dystroglycan and muscle-specific laminin) components (Ehmsen et al., 2002). More recently, dysregulation of dystrophin and other DAPC members has been observed in cancer (Acharyya et al., 2005), atrophy (Risson et al., 2009; Swiderski et al., 2021; Chockalingam et al., 2002), and aging (Hughes et al., 2017; Townsend et al., 2011; Hord et al., 2016; Kosek and Bamman, 2008), leading to decreased muscle integrity. Despite their importance, mechanisms underlying regulation of expression of *dystrophin* and other DAPC members are largely unknown.

Reversible acetylation of proteins is a well-known component of the regulation of gene expression in response to changes in the cellular environment. Lysine acetylation is catalyzed by lysine acetyl transferases (KATs), and can be removed by lysine deacetylases (KDACs). Importantly, dysregulation of KAT and KDAC activity occurs in various diseases including cancer (Gil et al., 2017), Alzheimer’s (Cohen et al., 2011), cardiovascular disease (Li et al., 2020), muscular dystrophy (Consalvi et al., 2011, 2013; Bettica et al., 2016), and aging. Post-translational modifications (PTMs) such as lysine acetylation influence secondary structure, inter-or intramolecular interactions, or catalytic activity. In particular, acetylation of lysine residues on transcription factors are a well characterized PTM that can affect binding to DNA or cofactors for the regulation of gene expression (Narita et al., 2019). In both muscle disease and aging inhibition of KDAC activity has been shown to result in beneficial effects on muscle (reviewed in (McIntyre et al., 2019)), however the mechanisms linking protein acetylation to muscle integrity are poorly understood.

As an evolutionarily conserved KAT, the lysine acetyltransferase action of the KAT protein general control of amino acid synthesis protein 5 (GCN5; KAT2A) is capable of acetylating both histone and non-histone lysines. Acetylation by GCN5 has been shown to stabilize the c-Myc oncoprotein (Patel et al., 2004), or recruit other HATs for the transcriptional co-activation of tumor suppressor p53 (Barlev et al., 2001), which in each case activate transcription. The acetylation of the metabolic coactivator PGC-1α in liver made the first direct link between GCN5 activity and the inhibition of PGC-1α-driven regulation of gluconeogenesis (Lerin et al., 2006). From these findings arose the hypothesis that GCN5 deletion could drive PGC-1α-directed mitochondrial biogenesis in muscle. However, recent work could not identify a metabolic role for GCN5 in skeletal muscle (Dent et al., 2017; Svensson et al., 2020).

To further interrogate the role of GCN5 acetylation in muscle, we performed an unbiased transcriptome analysis of the muscle-specific KO of GCN5 (*Gcn5*^skm-/-^) that best correlated to muscle atrophy and dystrophy datasets. Additionally, *Gcn5*^skm-/-^ mice exhibited a reduction in dystrophin expression that when interrogated *in vivo,* with different experimental models of eccentric muscle contraction, presented with a myopathic phenotype. To explain the regulation of dystrophin and other components of DAPC expression in we identified a role for GCN5 in acetylation of the DNA-binding zinc finger domain of the transcription factor/repressor ying yang 1 (YY1). We conclude that dystrophin and components of the DAPC are positively regulated by GCN5-directed acetylation of YY1.

## Results

### Transcriptomic analysis reveals the dysregulation of dystrophin and genes associated with the DAPC in Gcn5^skm-/-^ mice

We first set out to generate *Gcn5*^skm-/-^ mice in order to perform an unbiased transcriptome analysis that would inform of potential phenotypes and signalling pathways governed by GCN5 in differentiated muscle. To do this a conditional mouse knockout for GCN5 was generated by flanking exons 5 and 6 of *Gcn5* with LoxP sites. Conditional *Gcn5* knockout mice were then crossed with mice expressing *Cre* from a transgenic human *α-skeletal actin* (Hsa) gene promotor (Miniou et al., 1999) (Fig. S1), resulting in *Gcn5*^skm-/-^ mice (Fig. 1a). Note that *Gcn5*^skm-/-^ muscle tissue contains residual *Gcn5* mRNA from resident non-muscle cells, vasculature etc., as is typical of loss-of-function models generated using the HSA-*Cre* mouse line.

**Figure 1.**
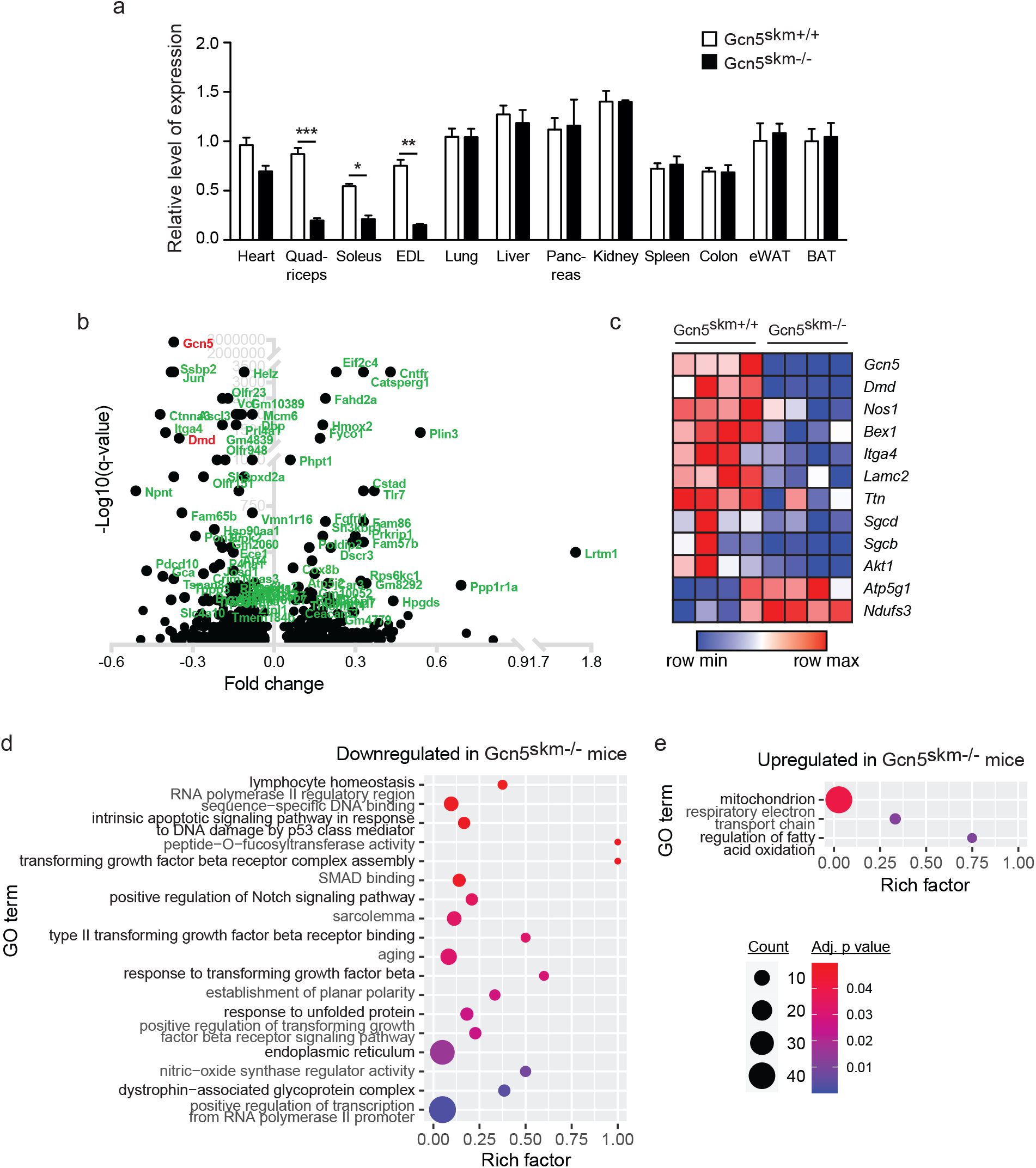
Validation and transcriptomic profiling of *Gcn5*^skm-/-^ mice. (a) Relative gene expression analysis of *Gcn5* in various tissues from control and *Gcn5*^skm-/-^ mice as determined by RT-qPCR. Values were normalized to 36B4. Data presented as mean +/-S.E.M. for controls and *Gcn5*^skm-/-^ mice. n = 10-11/group. \**p* < 0.05, \*\**p* < 0.01, \*\*\**p* < 0.001 versus *Gcn5*^skm+/+^ as measured by two-tailed Student’s *t* test. (b) Volcano plot of 1172 differentially expressed genes from *Gcn5*^skm-/-^ gastrocnemius muscle, representing the top 100 most significant genes. (c) Heat map showing dysregulated genes for DAPC and membrane proteins (Dmd, Sgcd, Sgcb, Nos1), ECM proteins (Lamc2, Itga4), energy metabolism proteins (Atp5g1, Ndufs3, Akt1) and Bex1, a marker of muscle regeneration, for control and *Gcn5*^skm-/-^ gastrocnemius muscle. n = 4/group. (d) Bubble plots showing the distribution and size of over-represented gene ontology pathways of the downregulated geneset, and (e) the upregulated geneset of gastrocnemius muscle from *Gcn5*^skm-/-^ mice. n = 4/group.

In order to further assess effects of loss of GCN5 in muscle, microarray analysis was performed on cDNA libraries generated from gastrocnemius muscle harvested from 6-month-old *Gcn5*^skm-/-^ mice and control littermates. cDNAs corresponding to a total of 1,236 microarray features were differentially expressed in the gastrocnemius muscle tissue of *Gcn5*^skm-/-^ mice, of which 789 were downregulated and 447 were upregulated (Supplementary excel file 1). Upon examination, dystrophin (*Dmd*) was among the most downregulated genes in *Gcn5*^skm-/-^ muscle (Fig. 1b), along with the dysregulation of other DAPC members, including, beta-sarcoglycan (*Sgcb*), delta-sarcoglycan (*Sgcd*), and the structural muscle protein transcript titin (*Ttn*) (Fig. 1c).

The InnateDB (Breuer et al., 2013) was used for gene ontology (GO) analysis of differentially expressed genes. Several pathways including dystrophin-associated glycoprotein complex, sarcolemma, nitric-oxide synthase (NOS) regulator activity, response to transforming growth factor beta were found to be downregulated in *Gcn5*^skm-/-^ muscle (Fig. 1d). Interestingly, in agreement with the hypothesis that *Gcn5* negatively regulates mitochondrial biogenesis through acetylation and inactivation of PGC-1α, and despite the absence of a metabolic phenotype in our study, mitochondrion respiratory electron transport chain and regulation of fatty acid oxidation pathways were upregulated in *Gcn5*^skm-/-^ muscle (Fig. 1e).

### GCN5 regulates dystrophin protein expression and maintains muscle integrity during eccentric contractions and with age

To confirm the reduction in *Dmd* gene expression observed using transcriptomics we performed both biochemical and histological analyses on *Gcn5*^skm+/+^ and *Gcn5*^skm-/-^ muscle samples. RT-qPCR on mRNA isolated from control and *Gcn5*^skm-/-^ gastrocnemius muscle showed decreased *Dmd* transcript (Fig. 2a). Immunostaining and western immunoblotting of dystrophin protein on tibialis anterior (TA) muscle samples, confirmed that dystrophin protein levels were reduced in *Gcn5*^skm-/-^ muscle compared to control mice (Fig. 2b-c). GCN5 regulation of dystrophin protein expression was also confirmed in C2C12 myotubes using adenoviral-driven shRNA constructs to knockdown *Gcn5* (Fig. 2d).

**Figure 2.**
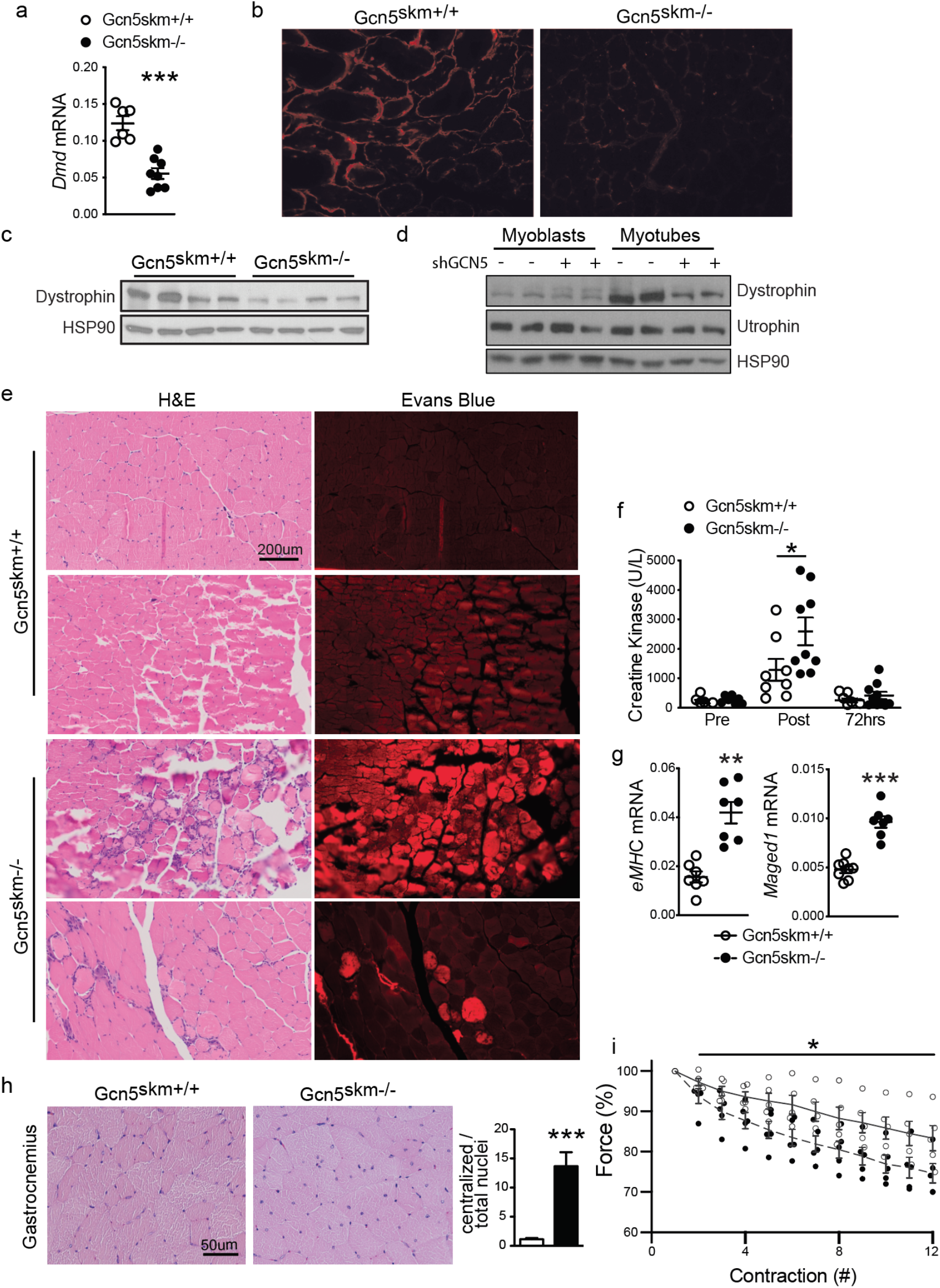
GCN5 regulates dystrophin protein expression and maintains muscle integrity during eccentric contractions and aging. (a) Relative expression of *Dmd* and *Utrn* in gastrocnemius muscle from control and *Gcn5*^skm-/-^ mice as measured by RT-qPCR relative to housekeeping genes *36b4* and *Gapdh*. Data presented as mean +/-S.E.M. for controls and *Gcn5*^skm-/-^ mice. n = 8-11/group. \*\**p* < 0.01 versus *Gcn5*^skm+/+^ as measured by two-tailed Student’s *t* test. (b) Representative images of immunostaining of TA muscle sections from control and *Gcn5*^skm-/-^ mice for dystrophin protein (red). (c) Western blot of whole protein extract from gastrocnemius muscle of control and *Gcn5*^skm-/-^ mice using anti-dystrophin with anti-HSP90 antibodies used as loading control. (d) Western blot of protein extract from C2C12 myoblasts and myotubes after 4 days of differentiation in the presence or absence of adenoviral-driven shGCN5 using anti-dystrophin and anti-utrophin antibodies with anti-HSP90 antibodies used as loading control. (e) Representative sections of TA muscle from control and *Gcn5*^skm-/-^ mice after downhill running stained with H&E (left column; brightfield) and Evans blue (right column; fluorescence). Two representative images for each genotype taken from separate mice. (f) Plasma creatine kinase levels in control and *Gcn5*^skm-/-^ mice measured before (Pre), directly after (Post) and 72 hours after (72hrs) downhill running. Data presented as mean +/-S.E.M. for controls and *Gcn5*^skm-/-^ mice. n = 7-9/group. * denotes *p* < 0.05 via ANOVA with Tukey’s post-hoc test versus *Gcn5*^skm+/+^. (g) Relative expression of embryonic myosin heavy chain (*eMHC*) and *Maged1* in control and *Gcn5*^skm-/-^ gastrocnemius tissue harvested 72 hours after downhill running. Data presented relative to housekeeping genes *36b4* and *Gapdh* and as mean +/-S.E.M. for controls and *Gcn5*^skm-/-^ mice. n = 7-8/group. \*\**p* < 0.01, \*\*\**p* < 0.001 versus *Gcn5*^skm+/+^ as measured by twotailed Student’s *t* test. (h) Representative sections of gastrocnemius muscle from 12-month-old control and *Gcn5*^skm-/-^ mice stained with H&E (brightfield). Number of centralized nuclei per total nuclei in control *Gcn5*^skm-/-^ mice were quantified using 3 fields per section. Data presented as mean +/-S.E.M. for controls and *Gcn5*^skm-/-^ mice. n = 3/group. *** denotes *p* < 0.001 versus *Gcn5*^skm+/+^ as measured by two-tailed Student’s *t* test. (i) Force measurement (percentage of initial contraction) of consecutive eccentric contractions of diaphragm strips harvested from control and *Gcn5*^skm-/-^ mice. Data presented as mean +/-S.E.M. for controls and *Gcn5*^skm-/-^ mice. n = 5/group. * denotes *p* < 0.05 via ANOVA versus *Gcn5*_skm+/+_.

Because *Gcn5*^skm-/-^ mice have decreased *Dmd* expression and exhibit a dystrophic gene expression profile, mice were further assessed for an underlying muscle disorder. *Gcn5*^skm-/-^ mice had no difference in percent lean mass, fat mass or relative muscle weight compared to controls (Fig. S2a-b). When mice were subjected to a moderate protocol of downhill running, a model that can result in muscle damage due to repetitive eccentric muscle contractions (EC) (Armstrong et al., 1983), TA muscles of *Gcn5*^skm-/-^ mice demonstrated higher levels of Evans dye uptake (Fig. 2e), indicating increased permeability of the muscle sarcolemma due to damage (Schmalbruch, 1975). There was no difference in distance ran between *Gcn5*^skm-/-^ mice and controls (Fig. S2c), however immediately following the downhill running protocol, circulating creatine kinase levels, an early biomarker of muscle damage (Bulfield et al., 1984), were further elevated in *Gcn5*^skm-/-^ mice compared to their controls (Fig. 2f), with no significant changes in creatinine, aspartate aminotransferase or total protein levels (Fig. S2d–f). Downhill running in *Gcn5*^skm-/-^ mice elicited increases in transcript levels for markers of muscle regeneration after 72hrs, including embryonic myosin heavy chain (*eMhc*) and *Maged1* (Fig. 2g), indicating that the damage caused by EC triggered a compensatory increase in muscle regeneration in *Gcn5*^skm-/-^ animals. Aged *Gcn5*^skm-/-^ muscle also exhibited a susceptibly to damage as evidenced by an increased proportion of centralized nuclei in the gastrocnemius and quadriceps muscles (Fig. 2h and S2g).

To further examine susceptibility to muscle damage and sarcolemma stability in *Gcn5*^skm-/-^ mice, diaphragm, a muscle that exhibits profound progressive pathology in dystrophic animals (Stedman et al., 1991; Addicks et al., 2018), was excised and mounted on a force transducer apparatus while being physiologically maintained *ex vivo* to measure loss of force during repeated eccentric contractions. *Ex-vivo* eccentric contraction of the diaphragm muscle allows real time assessment of muscle damage by quantifying muscle force during repeated eccentric contractions (Addicks et al., 2018). To ensure diaphragm strips were correctly prepared and were tolerant to repetitive non-eccentric contractions, muscles were maintained *in vitro* for 60 minutes and then subjected to repeated non-eccentric contractions for 12 minutes with no loss of force (Fig. S2h). Diaphragm strips from *Gcn5*^skm-/-^ mice exhibited a more rapid decline in muscle force during a series of eccentric contractions as compared to controls (Fig. 2i), further indicating an intrinsic susceptibility to damage caused by the loss of *Gcn5*.

### A transcriptome meta-analysis of Gcn5^skm-/-^ muscle revealed human muscle atrophy and muscular dystrophy as the top correlated disease biosets

One thousand two hundred thirty-six (1,236) differentially expressed cDNAs from *Gcn5*^skm-/-^ gastrocnemius muscles were mapped to 1171 genes in the Illumina BaseSpace™ Correlation Engine (Kupershmidt et al., 2010), and processed for further meta-analysis with a focus on highly correlated musculoskeletal diseases. The *Gcn5*^skm-/-^ muscle differentially expressed transcriptomic bioset was positively correlated with nine musculoskeletal diseases, among which muscle atrophy (MA) and Duchenne muscular dystrophy had the highest correlation scores (Fig. 3a). A targeted meta-analysis was performed between the *Gcn5*^skm-/-^ bioset and twelve individual significantly correlated biosets obtained from four MA studies (Mazzatti et al., 2008; Bialek et al., 2011; Jackman et al., 2012; Chua et al., 2015), and three significantly correlated biosets obtained from three DMD studies (Bakay et al., 2006; Bachinski et al., 2010; Khairallah et al., 2012). Differentially expressed genes for each study were obtained and correlated with the *Gcn5*^skm-/-^ bioset (Supplementary excel file 2). Among 741 downregulated genes in the *Gcn5*^skm-/-^ bioset, 314 (42.4%) were downregulated in MA, 236 (31.8%) were downregulated in DMD, and 118 (15.9%) were downregulated in both MA and DMD (Fig. 3b). Among 431 upregulated genes in the *Gcn5*^skm-/-^ bioset, 162 (37.6%) were upregulated in MA, 134 (31.1%) were upregulated in DMD, and 56 (13.0%) were upregulated in both MA and DMD.

**Figure 3.**
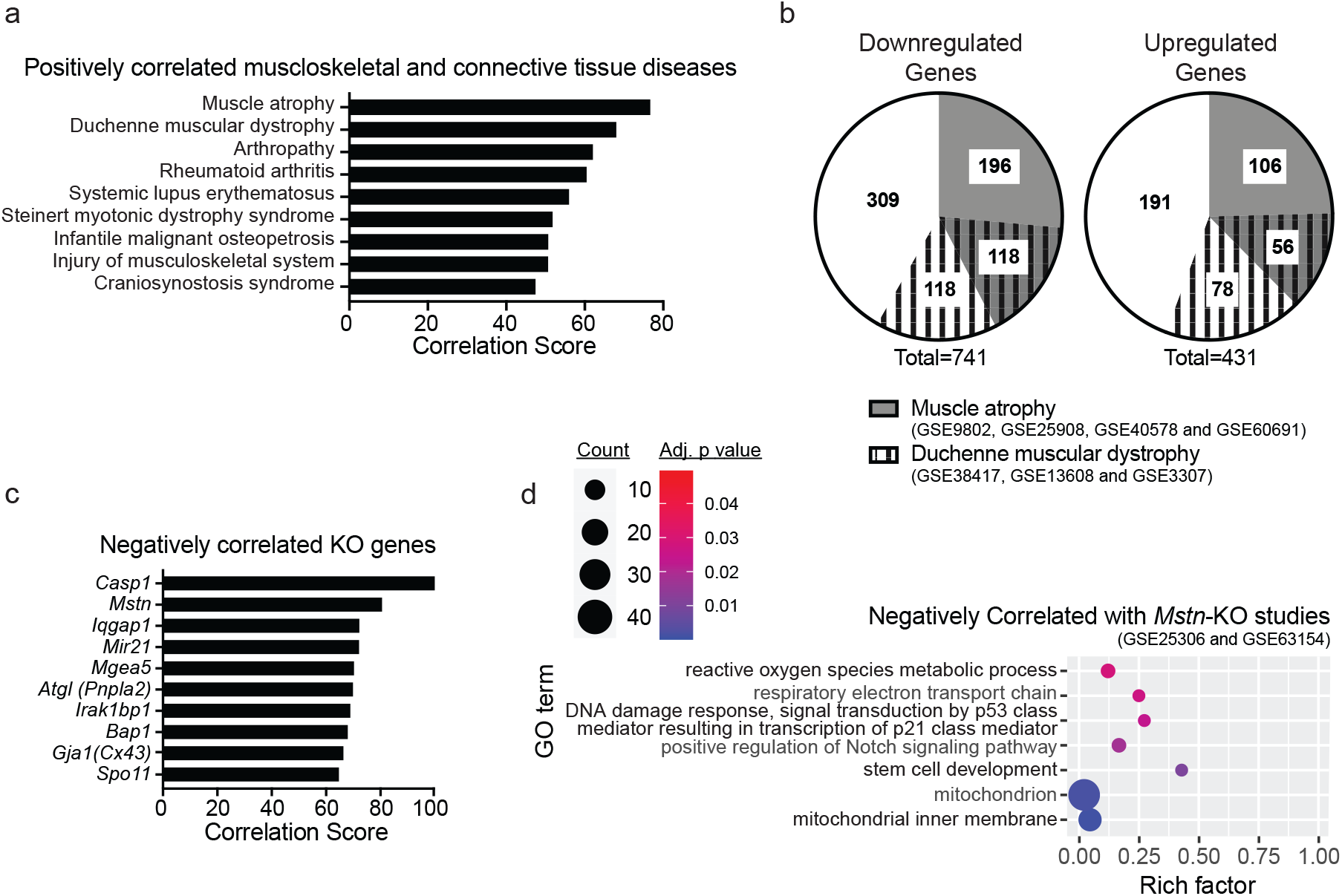
Dysregulation of *Dmd* and DAPC gene expression in *Gcn5*^skm-/-^ muscle is positively correlated with muscle atrophy and dystrophy. (a) Bar graph showing all positively correlated musculoskeletal diseases obtained from BaseSpace Correlation Engine by using all differentially expressed genes from the *Gcn5*^skm-/-^ bioset as an input for the meta-analysis. (b) Pie charts representing commonality among the *Gcn5*^skm-/-^, muscle atrophy and Duchenne muscular dystrophy biosets following a meta-analysis for downregulated and upregulated genes. (c) Bar graph showing negatively correlated KO genes obtained from BaseSpace Correlation Engine by using all differentially expressed genes from the *Gcn5*^skm-/-^ bioset as an input. (d) Bubble plots showing the distribution and size of negatively correlated over-represented gene ontology pathways between the *Gcn5*^skm-/-^ differentially expressed geneset and *Mstn-KO* genesets (GSE25306 and GSE63154).

Further supporting an atrophy phenotype, a similar meta-analysis was performed which specifically compared the *Gcn5*^skm-/-^ muscle bioset to all available mouse knockout biosets from any tissue (Fig. 3c and S3a). Of those identified the most negatively correlated bioset from muscle tissue was from a germline *Mstn-KO* (p-value < 0.0017), a mouse model that exhibits extensive muscle hypertrophy (Fig. 3c and Supplementary excel file 3). Of the *Gcn5*^skm-/-^ muscle bioset of 1171 up and downregulated genes, 226 (19.3%) were found to be negatively correlated with germline *Mstn*-KO muscle biosets (Rahimov et al., 2011; Yang et al., 2015) (Fig. S3b; Supplementary excel file 3). GO term analysis comparing the *Gcn5*^skm-/-^ bioset with *Mstn-KO* biosets was used to examine negatively and positively correlated pathways (Fig. 3d and S3c). Notably, the meta-analysis demonstrated a lower Notch signaling pathway geneset expression in the *Gcn5*^skm-/-^ bioset compared to the *Mstn*-KO biosets, a pathway known to be reduced in muscular dystrophy (Church et al., 2014; Vieira et al., 2015), and important for muscle regeneration (Bi et al., 2016). Several mitochondrial pathways were also negatively correlated, in agreement with increased mitochondrial pathways in *Gcn5*^skm-/-^ muscle when compared to the hypertrophic response in *Mstn*-KO muscle. In addition, a p53-mediated pathway was negatively correlated and downregulated in *Gcn5*^skm-/-^ muscle, which is supported by previous reports for GCN5-directed stabilization of p53. In contrast, the most positively correlated mouse knockout biosets (Fig. S2a), included the muscle specific sodium channel *Clcn1*, for which mutations in the gene is associated with myotonia (Pusch, 2002), the remaining positively correlated biosets were not derived from muscle tissue.

### YY1 represses Dmd and DAPC expression in mouse myotubes and is correlated to reduced structural gene expression and muscle fibre diameters in humans

In order to determine the mechanism by which GCN5 might regulate the expression of *Dmd* and other members of the DAPC, TRANSFAC (TRANScription FACtor database (Matys, 2006)), a manually curated database of eukaryotic transcription factors that includes their genomic binding sites and DNA binding profiles, was queried and YY1, a Gli-Kruppel zinc finger family protein, was identified as a probable factor to regulate *Dmd* gene expression (Supplementary excel file 4). Previous work showed that YY1 negatively regulates *Dmd* expression in muscle myoblasts cells (Galvagni et al., 1998; Zanotti et al., 2015). Similarly, YY1 was shown to negatively regulate dystrophin Dp71, the smallest protein encoded by the *Dmd* gene, in hepatic cells (Peñuelas-Urquides et al., 2016). Finally, YY1 appeared to be a potential target of GCN5 acetylation as YY1 was previously shown to be a target of acetyltransferases p300 and PCAF (Yao et al., 2001).

To determine if YY1 was a likely target of GCN5 control, we performed a meta-analysis using BaseSpace Correlation Engine between muscle biosets of *Gcn5*^skm-/-^ and muscle-specific *Yy1* KO mice (GSE39009) (Blättler et al., 2012b). This analysis provided a significant (p-value=0.0057) negative correlation with a total of 151 (12.9%) genes being negatively correlated (Fig. S4a, Supplementary excel file 5). GO term analysis through InnateDB (Breuer et al., 2013) identified dysregulated pathways that were positively (Fig. S4b) and negatively (Fig. 4a) correlated between the *Gcn5*^skm-/-^ and muscle-specific *Yy1* KO biosets (Blättler et al., 2012b). These included dystroglycan binding and nitric-oxide synthase regulator activity, both reduced genesets in the *Gcn5*^skm-/-^ versus *Gcn5*^skm+/+^ bioset. These results would imply that GCN5 is negatively correlated to YY1 with respect to directing changes in markers of muscular dystrophy.

**Figure 4.**
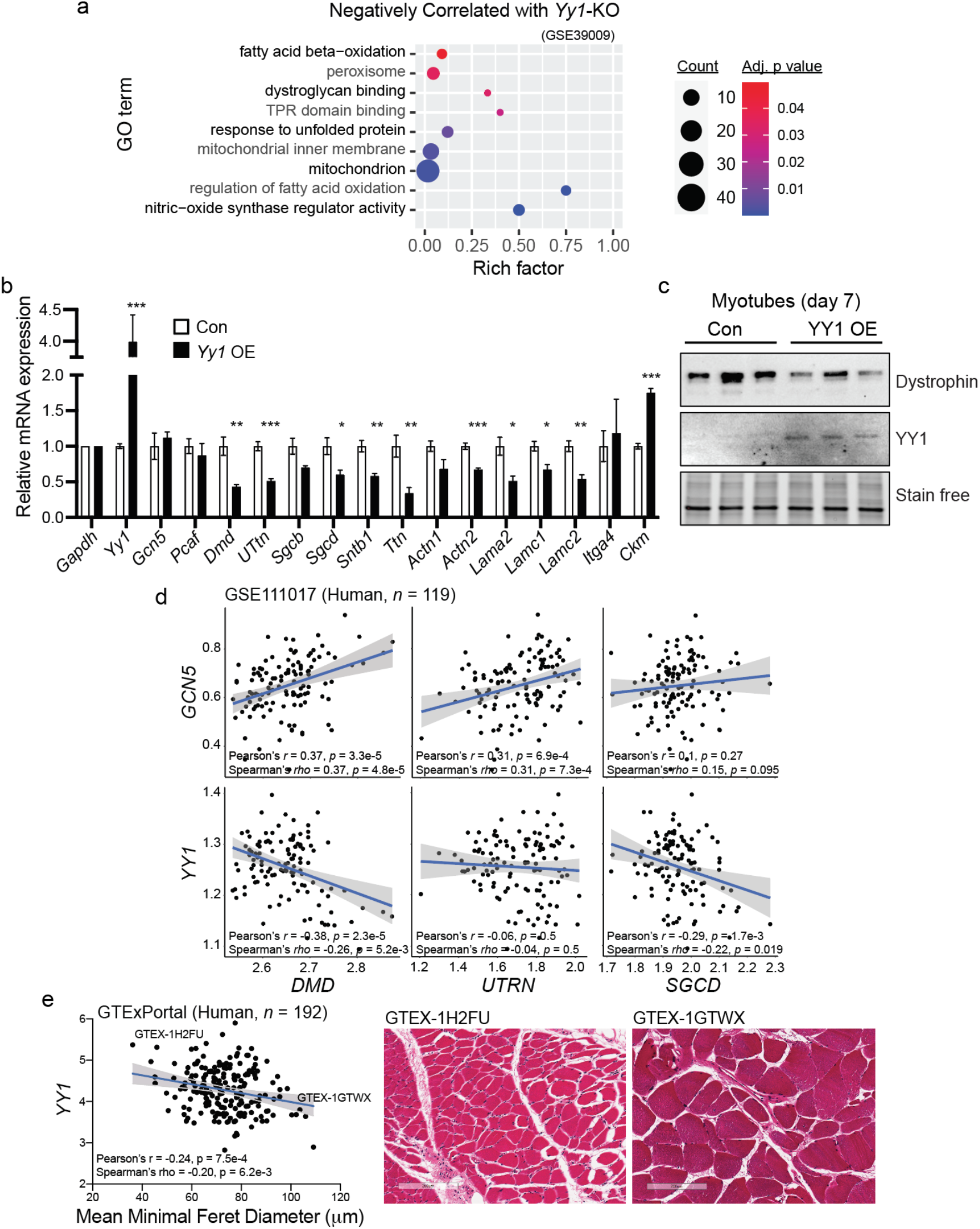
YY1 regulates dystrophin and DAPC expression and is negatively correlated to human fibre diameter. (a) Bubble plots showing the distribution and size of negatively correlated over-represented gene ontology pathways between the *Gcn5*^skm-/-^ differentially expressed geneset and a *Yy1-KO* study geneset (GSE39009). (b) Relative expression of genes in differentiated primary mouse myotubes from 4 individual empty vector control (Con) or *Yy1* overexpressing (*Yy1* OE) lentivirally transduced colonies for *Gcn5*, *Pcaf*, DAPC, cell surface, and ECM protein expressing genes (*Dmd*, *Utrn*, *Sgcb*, *Sgcd*, *Sntb1*, *Actn1*, *Actn2*, *Lama2*, *Lamc1*, *Lamc2*, *Itga4*) and muscle specific genes *Ttn* and *Ckm* as determined by RT-qPCR. n=4 (from 4 independent colonies selected after lentiviral transduction). \**p* < 0.05, \*\**p* < 0.01, \*\*\**p* < 0.001 versus *Gcn5*^skm+/+^ as measured by two-tailed Student’s *t* test. (c) Western blot of Dystrophin protein from empty vector control (Con) and YY1 overexpressing (YY1 OE) mouse primary myotubes (top), and stain free protein loading control (bottom). (d) Scatter plots showing correlations of either *GCN5* or *YY1 transcripts* (Z-score of TMM-normalized TPM) with three representative structural muscle transcripts (*DMD*, *UTRN*, and *SGCD*) in human skeletal muscle transcriptome (GSE111017, n = 119). (e) Correlation between mean minimum feret diameter of human muscle fibers, consisting of 250 measured muscle fibers from 192 GTEx samples (GTExPortal database v8), and *YY1* (Log2 TPM) transcript expression. Representative images of GTEx muscle samples are shown.

To investigate the potential negative relationship between GCN5 and YY1 on the expression of *dmd* and other DAPC members, YY1 was overexpressed in primary mouse myotubes in cell culture and gene expression was assessed through qPCR (Fig. 4b). Notably, several genes that were downregulated in *Gcn5*^skm-/-^ muscle, including *Dmd, Sgcb, Sgcd* and *Ttn* (Fig. 1c), were also downregulated by exogenous YY1 in primary myotubes. Overexpression of YY1 did not affect expression of GCN5 or PCAF or muscle specific creatine kinase (*Ckm*). Western blotting also confirmed that dystrophin expression is decreased in primary mouse myotubes with exogenous YY1 expression (Fig. 4c), which is complemented by our observed reduction in myotube dystrophin expression with *Gcn5* knockdown (Fig. 2d).

In human muscle transcriptome (GSE111017, n = 119), the correlations between transcripts for muscle structural proteins (i.e., *DMD*, *UTRN*, and *SGCD*) and either *GCN5* or *YY1* were compared. *GCN5* exhibits positive correlations with three representative transcripts, *DMD*, *UTRN*, and *SGCD*, although *SGCD* does not reach statistical significance. Conversely, *YY1* shows negative correlations with three genes, but *UTRN* did not reach significance (Fig. 4d). Finally, corroborating the positive correlation between *Gcn5*^Skm-/-^ muscle and muscle atrophy biosets (Fig. 3a and 3b), in human muscle (GTExPortal database v8; n=192) *YY1* transcript expression was negatively correlated to the mean minimum feret diameter of muscle fibers (Fig. 4e). This dataset also showed a positive correlation between *GCN5* and *DMD* transcript expression (Fig. S4c). These human transcriptomic and phenotype correlations drove us to hypothesize that GCN5 may acetylate and inhibit YY1 to maintain *dmd* and DAPC component expression for the maintenance of muscle integrity.

### YY1 is an acetylation target of GCN5

Given that YY1 is known to be acetylated by p300 and PCAF (Yao et al., 2001), we predicted that YY1 may be a target for GCN5 acetylation and regulation. To determine if GCN5 affects YY1 acetylation, YY1 was immuno-precipitated from muscle extracts and probed with anti-acetyl lysine antibody, which revealed decreased acetylation of YY1 protein in *Gcn5*^skm-/-^ extracts compared to *Gcn5*^skm+/+^ controls (Fig. 5a). This result was further explored *in vitro* by creating GST-labeled YY1 fragments (F1, aa 1-190; F2, aa 180-300; F3, aa 300-414) (Fig. 5b) to help determine the relative location of GCN5-directed YY1 acetylation. *In vitro* acetylation of YY1 fragments with GCN5 identified acetylation sites on the C-terminal region of YY1, which includes zinc-finger motifs of the YY1 DNA-binding domain (F3, aa 300-414) (Fig. 5c). Using nanoscale liquid chromatography coupled to tandem mass spectrometry (nano-LC-MS/MS) of *in vitro* GCN5 acetylated YY1 two novel acetylation sites (K392 and K393) were identified (Fig. S5). A 3D structural analysis of the YY1 DNA-binding domain revealed that K392 and K393 reside in an unstructured region of zinc finger 4 outside of the zinc binding site (Fig. 5d-i) (Houbaviy et al., 1996). This model predicts that the positively charged lysine residues of K392/3 interact with the negatively charged phosphate backbone of bound DNA, thereby stabilizing DNA binding in a similar manner to lysine residues found in histone proteins. As a result, in this model, neutralization of the positive charge of K392/3 through acetylation could destabilize binding of YY1 to DNA.

**Figure 5.**
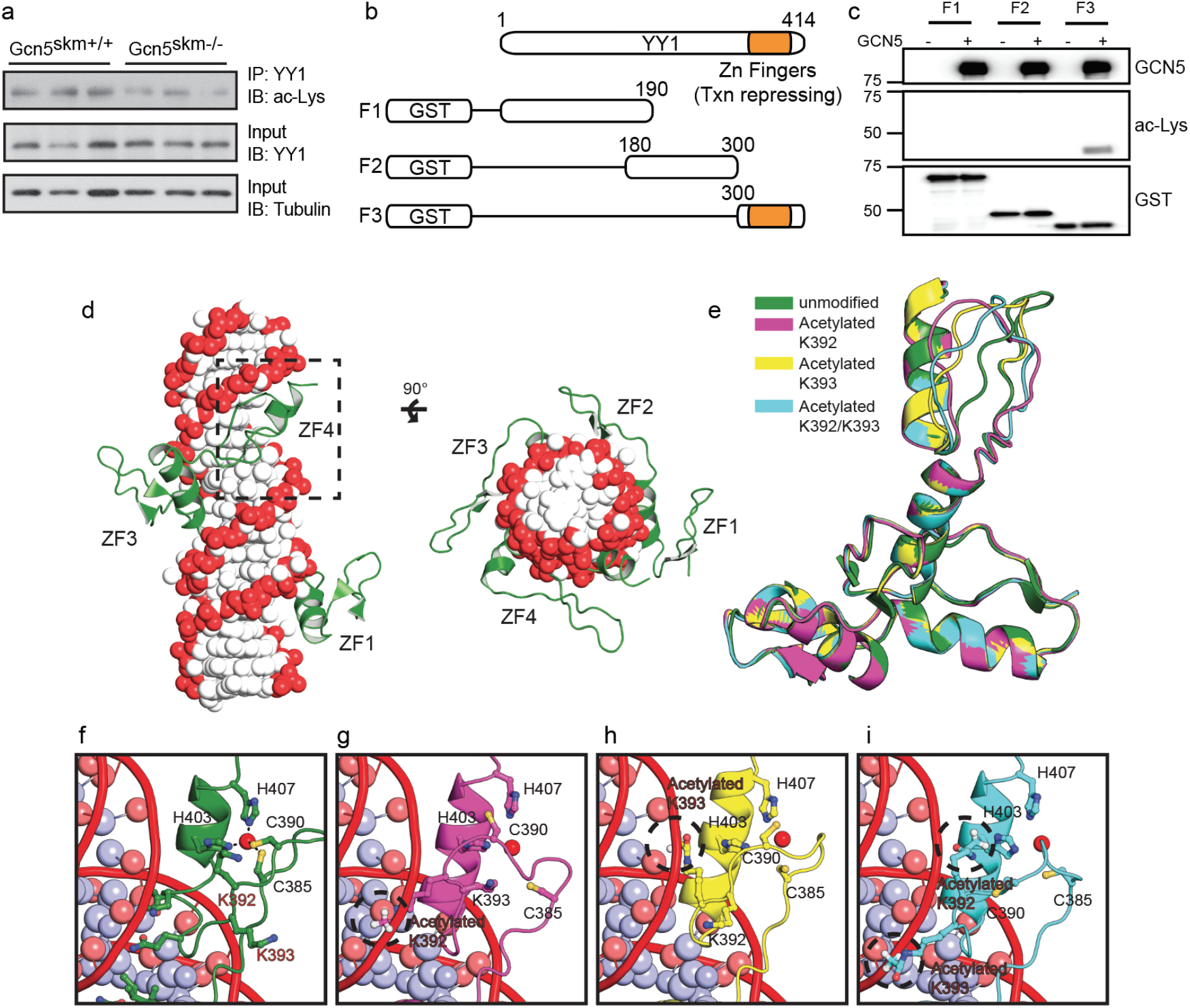
GCN5 acetylates YY1 and alters a predicted model of YY1 binding to DNA. (a) Western blot of anti-YY1 immunoprecipitated from controls and *Gcn5*^skm-/-^ mice probed with anti-acetyl-lysine (ac-Lys, top). Control blots for IP YY1 probed with anti-YY1 (middle) and total protein input for IP probed with anti-tubulin (bottom). (b) Schematic showing GST-tagged truncated YY1 used for *in vitro* acetylation (Figure 6C) and mass spectrometric analysis. (c) *In vitro* acetylation assay of the *in vitro* synthesized GST-YY1 fragments incubated with GCN5. YY1 fragments were incubated alone or with GCN5 (top). Acetylation of YY1-fragment (300-414) detected with anti-acetyl lysine (ac-Lys, middle). Input YY1 fragments detected by anti-GST antibody (bottom). (d) Crystal structure of YY1 zinc-finger region bound to DNA. (e-i) Modeling of disrupted YY1 zinc-finger structure and interactions with DNA when acetylated at acK392 and acK393.

### Acetylation of YY1 at K392 and K393 inhibits DNA binding and repression of Dmd

Through the use of *in vitro* acetylation assays of truncated YY1 fragments, p300 was shown to acetylate YY1 at a site between amino acids 170 and 200 (Yao et al., 2001). This same work found that PCAF acetylates YY1 at two residues located between amino acids 261 and 414 (Yao et al., 2001), however the precise location of acetylation by PCAF was not determined. To expand on our nano-LC-MS/MS analysis and confirm that K392 and K393 are the main acetylation sites of GCN5, both lysines were, independently and in tandem, converted to arginines through site-directed mutagenesis and subjected to *in vitro* acetylation by GCN5. Mutation of individual lysines resulted in decreased acetylation compared to the WT fragment, while mutation of both lysines resulted in a complete loss of acetylation *in vitro* (Fig. 6a).

**Figure 6.**
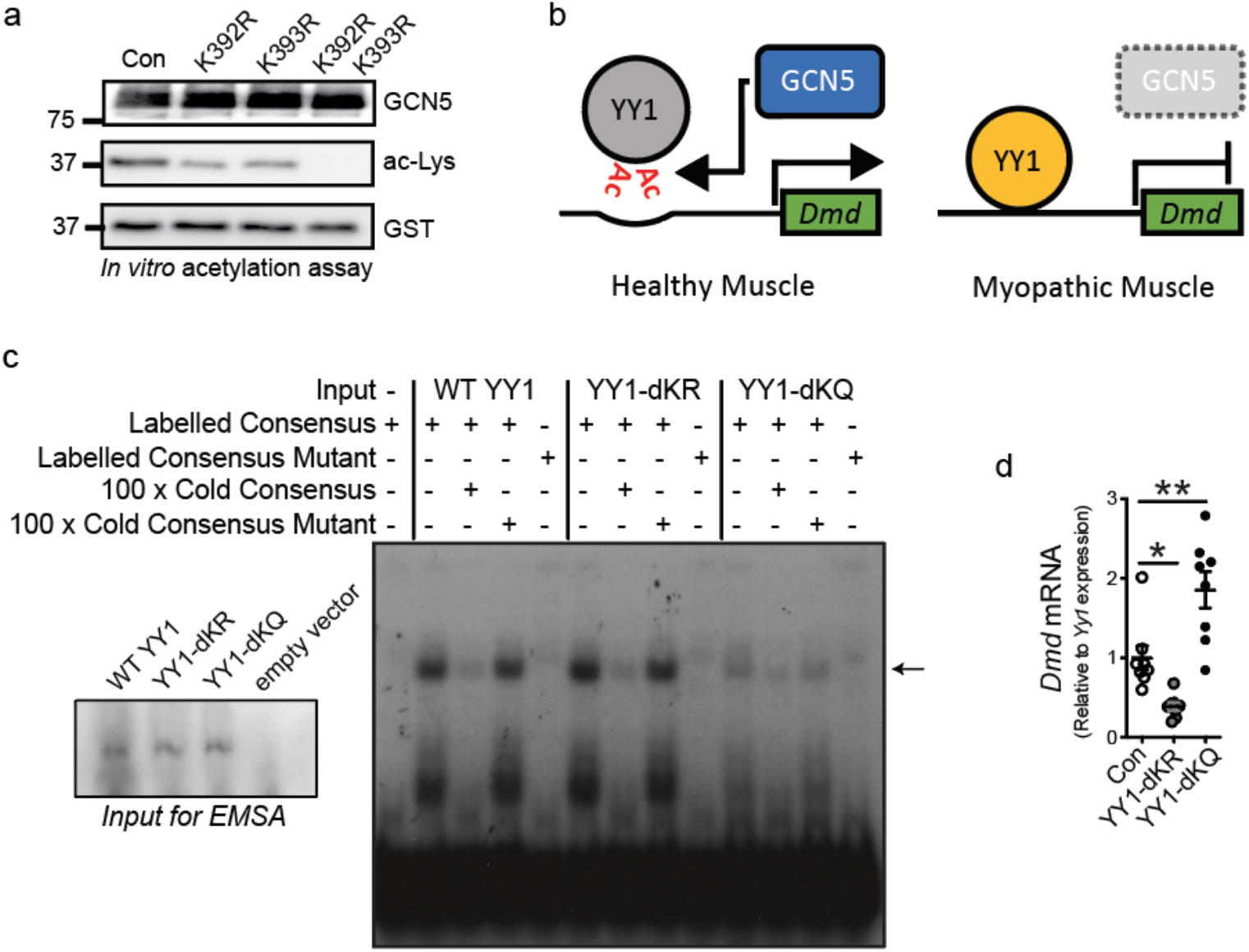
Acetylation suppresses YY1 DNA binding and inhibition of *Dmd* gene expression. (a) Western blot of *in vitro* acetylation by GCN5 of WT, YY1 (Con) or YY1 with lysines 292 and 293 mutated to arginine individually (K392R or K393R) or together (K392R/K393R) probed with anti-acetyl-lysine (ac-Lys). (b) Mechanism for the regulation of YY1 activity by GNC5. Under normal conditions GCN5 negatively regulates YY1 through acetylation and YY1 does not repress *Dmd* gene expression or other targets. In the absence of GCN5 regulation, YY1 abnormally represses *Dmd* and other genes resulting in a dystrophic phenotype. (c) Electrophoretic mobility shift assay (EMSA) of *in vitro* synthesized control YY1 (Con) or YY1 with lysines 392 and 393 mutated to arginine (YY1-dKR) or glutamine (YY1-dKQ). YY1 or YY1 mutants were incubated with radioactively labelled oligonucleotide dimers with consensus YY1 binding site (YY1 Consensus Labelled, lanes 2, 6 and 10), radioactively labelled YY1 consensus oligonucleotide dimers with 100-fold excess unlabelled YY1 consensus oligonucleotide dimers (100 X Cold Consensus, lanes 3, 7 and 11) or with 100-fold excess unlabelled mutated YY1 consensus oligonucleotide dimers (100 X Cold Mutant, lanes 4, 8 and 12), or with labeled mutated YY1 consensus oligonucleotide dimers (YY1 Mutant Labelled, lanes 5, 9 and 13). As a negative control for YY1 consensus binding, transcription-translation reaction without DNA was incubated with radioactively labelled YY1 consensus oligonucleotide dimers (lane 1). Bound oligonucleotide marked by arrow. Western blot of *in vitro* transcribed WT YY1, YY1 392/392 mutants and empty vector control in inset (lower left) showing uniform expression of YY1 control and mutant proteins used for EMSAs. (d) *Dmd* mRNA expression in C2C12 myotubes after 4 days of differentiation and transfected for 48hrs with FLAG-tagged YY1, YY1-dKR (K392R and K393R) or FLAG-YY1-dKQ (K392Q and K393Q). Data presented as mean +/-S.E.M. n = 8/group. \**p* < 0.05, \*\**p* < 0.01, \*\*\**p* < 0.001 via ANOVA with Tukey’s post-hoc test versus YY1 control.

Because the modeled YY1-DNA structure suggests that neutralization of positive charges on K392 and K393 through acetylation may result in inhibition of DNA binding by YY1, we postulated that GCN5-directed acetylation of YY1 destabilizes the YY1-directed inhibition of *Dmd* expression (Fig. 6b). To test whether acetylation of YY1 affects DNA binding, *in vitro* transcribed full-length WT YY1 and full-length YY1 mutants, that include K392 and K393 tandem mutations that mimic varied acetylation states, were used in electrophoretic mobility shift assays (EMSA) with labeled YY1-binding consensus oligonucleotide dimers. In agreement with our hypothesis, mutating both K392 and K393 to the uncharged acetylation mimic glutamine (YY1-dKQ) resulted in a loss of DNA binding, while mutating both lysines to the negatively charged arginine (YY1-dKR) retained DNA binding activity (Fig. 6c). To further establish a role for YY1 in the regulation of *Dmd* expression, WT YY1 and YY1 mutants were exogenously expressed in C2C12 myoblasts and differentiated for 48 hours after which cells were collected for RT-qPCR analysis. In alignment with our EMSA results, the YY1-dKR mutant, which cannot be acetylated but maintains the negative charge of the WT lysine residues resulted in decreased *Dmd* expression (Fig. 6d). The YY1-dKQ mutant, which does not bind DNA, resulted in increased *Dmd* expression likely through a dominant negative effect.

## Discussion

To examine the physiological role of GCN5 in postmitotic skeletal muscle, we crossed *Gcn5*^L2/L2^ mice with mice expressing *Cre* recombinase driven by a transgenic human *α-skeletal actin* gene promotor. Initial transcriptomic analysis of the resulting *Gcn5*^skm-/-^ mice identified reductions in the expression of dystrophin and other DAPC members. Similar to mouse models of muscular dystrophy, such as *mdx* mice that lack dystrophin expression, GCN5 deficiency in skeletal muscle leads to a progressive muscle myopathy that is primarily identified by a susceptibility to eccentric muscle damage.

While GCN5 is recognized as a general regulator of transcription, loss of GCN5 did not appear to have an effect on global transcription, likely due to redundancy with other histone acetyltransferases (Dent et al., 2017). An unbiased analysis of our transcriptomic dataset found multiple downregulated GO terms related to muscular dystrophy and muscle atrophy and aging. This was then further illustrated by a meta-analysis using BaseSpace Correlation Engine that determined that *Gcn5*^Skm-/-^ muscle best correlated to biosets of human muscle atrophy and Duchenne muscular dystrophy, when comparing our bioset to all publicly available data categorized by musculoskeletal and connective tissue diseases. *Gcn5*^Skm-/-^ muscle exhibited reduced expression of several critical muscle specific gene transcripts, with the most striking reduction being that of dystrophin. The links between *Gcn5*^Skm-/-^ and muscular disease were further emphasized by observed negative correlations between the *Gcn5*^Skm-/-^ bioset and that of KO mice for *Mstn, Cx43* and *Mir-21*, genes that have been linked to playing a negative role on *mdx* mouse phenotypes (Fig. 3c) (Nouet et al., 2020; Zanotti et al., 2015; Ardite et al., 2012; Dumonceaux et al., 2010). The finding that the *Gcn5*^Skm-/-^ bioset is negatively correlated with several *Mstn* KO biosets compliments findings that suggest myostatin pathway inhibition as a potential DMD therapeutic strategy (Campbell et al., 2017). Concomitant with gene expression changes, *Gcn5*^Skm-/-^ muscle also displayed a myopathic phenotype. *Gcn5*^Skm-/-^ muscle demonstrated increased fragility with eccentric contraction as evidenced by release of muscle specific creatine kinase into the circulation, uptake of Evans blue dye into the muscle from the circulation, a histological muscle damage phenotype and upregulation of regenerative gene expression, as is also seen in muscle pathology. Using a gold-standard assay to assess phenotype in mouse models of muscular dystrophy where muscle integrity is perturbed, we discovered a reduction in diaphragm contraction force produced during repeated eccentric contractions in *Gcn5*^Skm-/-^ muscle, confirming a hallmark phenotype of *mdx* mice (Addicks et al., 2018). The data therefore shows that GCN5 modulates dystrophin expression and maintains muscle integrity.

Because *Dmd* was among the most strongly downregulated genes in *Gcn5*^Skm-/-^ muscle we next aimed to determine how GCN5 might modulate *Dmd* expression and focused on YY1, a known regulator of *Dmd* (Galvagni et al., 1998). YY1 has been implicated as both a transcriptional repressor, as part of polycomb repressive complex 2, and as a transcriptional activator in other contexts (Shi et al., 1997). The *Yy1* knockout results in embryonic lethality at embryonic day 10.5 due to failure to properly form mesoderm (Donohoe et al., 1999). The muscle specific knockout of *Yy1* results in hyper-activation of insulin/IGF signaling, increased insulin/IGF pathway gene expression, and defective mitochondrial morphology and oxidative function (Blättler et al., 2012b; a). Upon further analysis of our *Gcn5*^Skm-/-^ muscle geneset, the DAPC-constituent-dystroglycan-binding GO term was found to be negatively correlated between genes that are differentially expressed between *Yy1* knockout muscle and control (Blättler et al., 2012b), and genes that are differentially expressed between *Gcn5*^Skm-/-^ muscle and control. Supporting this association, overexpression of YY1 in myotubes also resulted in decreased expression of *Dmd* and other DAPC members. Importantly, it was previously found that YY1 is acetylated by both p300 and the GCN5 homologue PCAF, and that PCAF based acetylation resulted in decreased DNA binding of YY1 (Yao et al., 2001). We also found that YY1 acetylation was strongly decreased in *Gcn5*^Skm-/-^ muscle suggesting that YY1 is acetylated by GCN5 *in vivo* and may modulate DNA binding. We then demonstrated that *YY1* gene expression was negatively correlated to muscle fibre size in data from the human GTEx consortium, indicating that YY1 may have a negative impact on muscle integrity in humans, as we have shown phenotypically in *Gcn5*^Skm-/-^ mice.

To specifically address whether GCN5 regulates the YY1 via acetylation, we performed an *in vitro* analysis that demonstrated GCN5-directed acetylation of YY1 at K392 and K393, which are known to locate within the C-terminal zinc finger of four zinc finger DNA binding sites on YY1 (Houbaviy et al., 1996). Indeed, EMSAs performed using control YY1 and YY1 mutants, mimicking acetylated and unacetylated K392/K393, revealed that mutation of the negatively charged lysine residues to uncharged glycine-acetylation-mimics resulted in a strong disruption of DNA binding; mutation to negatively charged arginine, which is not acetylated by GCN5 and mimics unacetylated lysine, resulted in stronger DNA binding. This supports the conclusion that acetylation of YY1 by GCN5 at K392/K393 disrupts DNA binding. Indeed, when YY1 mutants were expressed in myotubes, mutation of K392/K393 to glycine resulted in increased *Dmd* transcript expression, while mutation to arginine resulted in repression of *Dmd* expression.

KDAC inhibitors have been found to have beneficial effects for several types of muscle disorders (Minetti et al., 2006; Colussi et al., 2008; Consalvi et al., 2011; Johnson et al., 2013; Pambianco et al., 2016), and has led to Phase 2 and Phase 3 clinical trials that are currently underway for several forms of muscular dystrophy (Consalvi et al., 2013; Bettica et al., 2016). Our findings might suggest that the YY1 acetylation state in muscle could be maintained by HDAC inhibition to support muscle integrity, as a potential additional route of treatment effectiveness. These findings therefore advocate that considerable opportunity remains for a better understanding of the dynamic effects of protein acetylation in the regulation of muscle integrity for the potential treatment of muscle wasting diseases.

This work identifies a novel role for GCN5 as a regulator of gene expression through acetylation of YY1. As YY1 is a transcriptional regulator (Thomas and Seto, 1999; Meliala et al., 2020), acetylation and repression of YY1 DNA binding may also act to complement the known role of GCN5 in positive regulation of gene expression through acetylation of histones (Grant et al., 1997). Importantly, loss of GCN5 and its repressive acetylation of YY1 at K392/K393 results in YY1 based repression of genes of which have critical roles in muscle, most importantly dystrophin, resulting in an overall phenotype that resembles muscular dystrophy. These findings may be useful for the discovery of new therapeutics for maintaining healthy muscle during muscular dystrophy and other diseases.

## Acknowledgements

The authors are grateful for the technical support of Professor Bernard Jasmin and for resources at the Centre for Neuromuscular Disease, University of Ottawa, especially John Lunde from the facility. The authors are also thankful to Dr. James Flynn from Illumina for advice on *in silico* analysis.

## Author contributions

Conception of hypotheses: GA, GV, HZ, DR, JA and KJM. Design of the work: GA, HZ, DR, GV, PM, JA and KJM. Acquisition of the data: GA, HZ, DR, GV, PM, AG, SP, BK, DK, EK and KJM.

Drafting the manuscript: GA, GV, KJM. Analysis and interpretation of the data: GA, HZ, DR, GV, PM, AG, BK, DK, EW, JR, JA and KJM. All Authors: approved the final version of the manuscript.

## Supplementary Materials

**Figures S1 to S5 and Methods**

**Supplemental Excel File 1.** Microarray features and analysis from control and *Gcn5*^skm-/-^ mice. Related to Figures 1d and 1e.

**Supplemental Excel File 2.** Differentially expressed genes and analysis from studies related to Figure 3b.

**Supplemental Excel File 3.** Differentially expressed genes and analysis from studies related to Figures 3c, S3a and S3b.

**Supplemental Excel File 4.** TRANSFAC query of factors regulating *Dmd*

**Supplemental Excel File 5.** Differentially expressed genes and analysis from studies related to Figure S4a.

