## Supplemental Figures and Methods for "GCN5 Maintains Muscle Integrity by Acetylating YY1 to Promote Dystrophin Expression"

### Supplementary Figures and Methods

Fig S1.

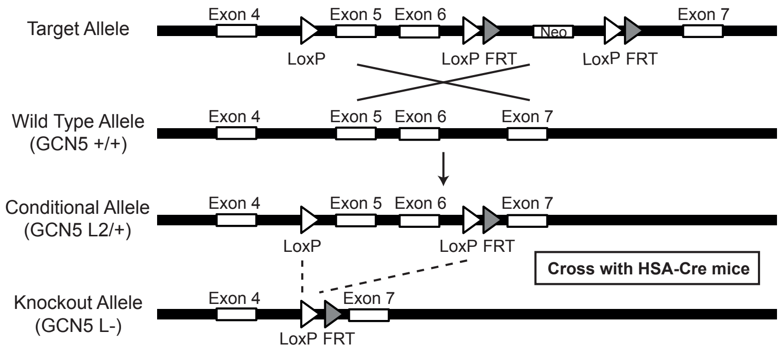
Target Allele
Exon 4
Exon 5
Exon 6
Neo
Exon 7
LoxP
LoxP FRT
LoxP FRT
Wild Type Allele
(GCN5 +/+)
Exon 4
Exon 5
Exon 6
Exon 7
Conditional Allele
(GCN5 L2/+)
Exon 4
Exon 5
Exon 6
Exon 7
LoxP
LoxP FRT
Cross with HSA-Cre mice
Knockout Allele
(GCN5 L-)
Exon 4
Exon 7
LoxP FRT

**Figure S1.** Generation of *Gcn5*$^{skm-/-}$ mice

Schematic of *Gcn5* gene targeting and conditional deletion of exons 5 and 6. Maps of the *Gcn5* genomic locus, the floxed allele with the neomycin cassette (+neo) (target allele) and without the neomycin cassette (−neo) (conditional allele). Positions of the exons are indicated.

Fig S2.

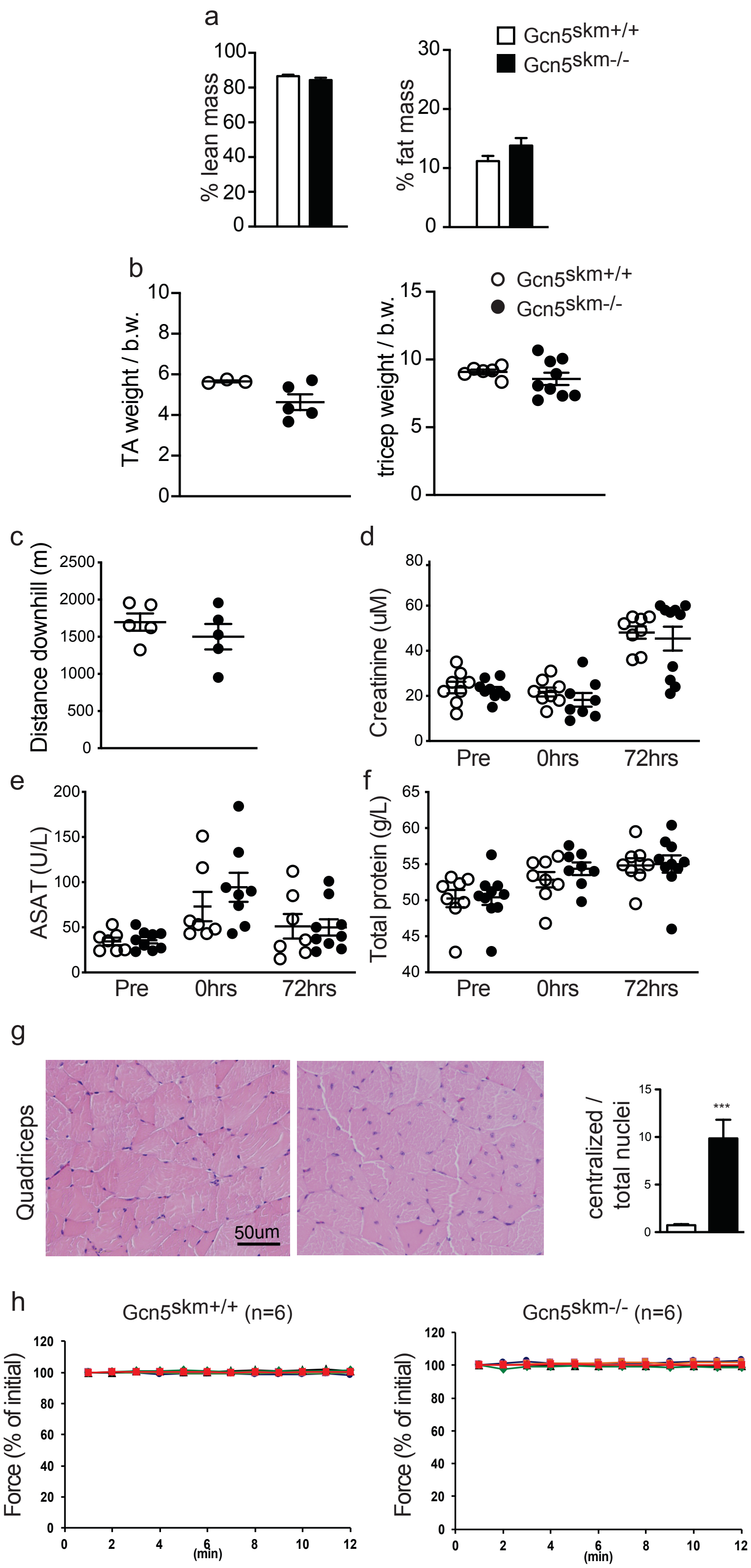
a
% lean mass
% fat mass
Gcn5skm+/+
Gcn5skm-/-
b
TA weight / b.w.
tricep weight / b.w.
c
Distance downhill (m)
d
Creatinine (uM)
e
ASAT (U/L)
f
Total protein (g/L)
Pre
0hrs
72hrs
g
Quadriceps
50um
centralized / total nuclei
***
h
Gcn5skm+/+ (n=6)
Gcn5skm-/- (n=6)
Force (% of initial)
(min)

**Figure S2.** GCN5 regulates dystrophin protein expression and maintains muscle integrity during eccentric contractions.

(a) Percent lean mass and fat mass for *Gcn5*$^{skm+/+}$ and *Gcn5*$^{skm-/-}$. Data presented as mean +/– S.E.M. for controls and *Gcn5*$^{skm-/-}$ mice. n = 12/group.

(b) Tibialis Anterior and triceps weights per amount of body weight for *Gcn5*$^{skm+/+}$ and *Gcn5*$^{skm-/-}$. Data presented as mean +/– S.E.M. for controls and *Gcn5*$^{skm-/-}$ mice. n = 3-8/group.

(c) Distance ran during downhill running for *Gcn5*$^{skm+/+}$ and *Gcn5*$^{skm-/-}$. Data presented as mean +/– S.E.M. for controls and *Gcn5*$^{skm-/-}$ mice. n = 5/group.

(d) Blood creatinine levels in control and *Gcn5*$^{skm-/-}$ mice immediately before (Pre), immediately after (0hrs) and 72 hours after (72hrs) downhill run. Data presented as mean +/– S.E.M. for controls and *Gcn5*$^{skm-/-}$ mice. n = 7-9/group.

(e) Blood ASAT levels in units per litre in control and *Gcn5*$^{skm-/-}$ mice immediately before (Pre), immediately after (0hrs) and 72 hours after (72hrs) downhill run. Data presented as mean +/– S.E.M. for controls and *Gcn5*$^{skm-/-}$ mice. n = 7-9/group.

(f) Total blood protein in grams per litre in control and *Gcn5*$^{skm-/-}$ mice immediately before (Pre), immediately after (0hrs) and 72 hours after (72hrs) downhill run. Data presented as mean +/– S.E.M. for controls and *Gcn5*$^{skm-/-}$ mice. n = 7-9/group.

(g) Representative sections of quadriceps muscle from 12-month-old control and *Gcn5*$^{skm-/-}$ mice stained with H&E (brightfield). Number of centralized nuclei per total nuclei in control *Gcn5*$^{skm-/-}$ mice were quantified using 3 fields per section. Data presented as mean +/– S.E.M. for controls and *Gcn5*$^{skm-/-}$ mice. n = 3/group. *** denotes $p < 0.001$ versus *Gcn5*$^{skm+/+}$ as measured by two-tailed Student's *t* test.

(h) Force produced during a series of contractions after 1 hour of equilibrium in apparatus and prior to initiation of eccentric contraction series for diaphragm strips used in figure 4D. Data presented as mean +/– S.E.M. for controls and *Gcn5*$^{skm-/-}$ mice. n = 5/group.

Fig S3.

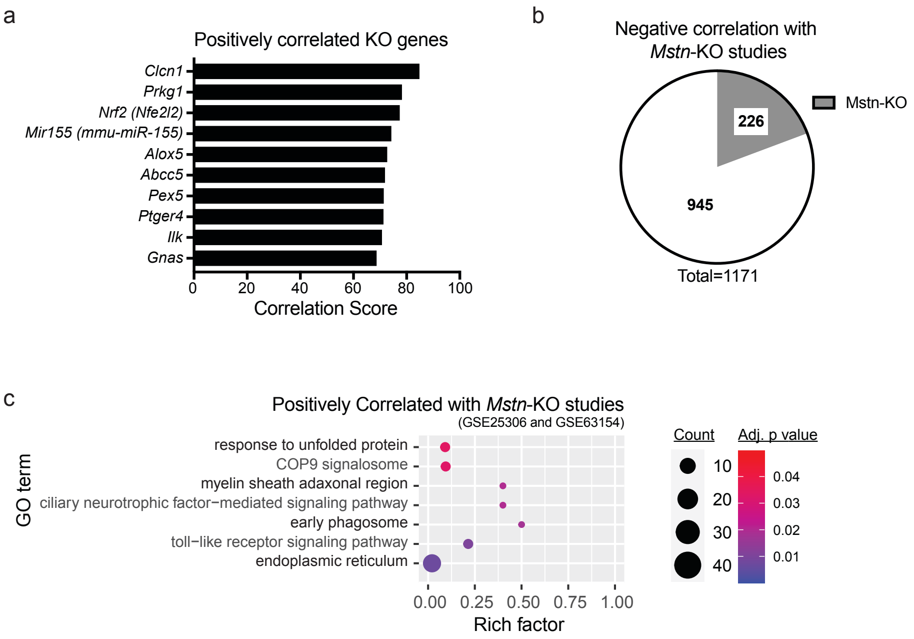
a
Positively correlated KO genes
Clcn1
Prkg1
Nrf2 (Nfe2l2)
Mir155 (mmu-miR-155)
Alox5
Abcc5
Pex5
Ptger4
Ilk
Gnas
0 20 40 60 80 100
Correlation Score
b
Negative correlation with Mstn-KO studies
226
945
Mstn-KO
Total=1171
c
Positively Correlated with Mstn-KO studies
(GSE25306 and GSE63154)
GO term
response to unfolded protein
COP9 signalosome
myelin sheath adaxonal region
ciliary neurotrophic factor−mediated signaling pathway
early phagosome
toll−like receptor signaling pathway
endoplasmic reticulum
0.00 0.25 0.50 0.75 1.00
Rich factor
Count
10
20
30
40
Adj. p value
0.04
0.03
0.02
0.01

**Figure S3.** Dysregulation of *Dmd* and DAPC gene expression in *Gcn5*$^{skm-/-}$ muscle is positively correlated with muscle atrophy or dystrophy.

(a) Bar graph showing negatively correlated KO genes obtained from Correlation Engine by using all differentially expressed genes from the *Gcn5*$^{skm-/-}$ bioset as an input.

(b) Pie chart representing negatively correlated genes between *Gcn5*$^{skm-/-}$ and *Mstn*-KO (GSE25306 and GSE63154) biosets following meta-analysis of the selected studies for downregulated and upregulated genes.

(c) Bubble plot showing the distribution and size of positively correlated over-represented gene ontology pathways between the *Gcn5*$^{skm-/-}$ differentially expressed bioset and *Mstn*-KO study biosets (GSE25306 and GSE63154).

Fig S4.

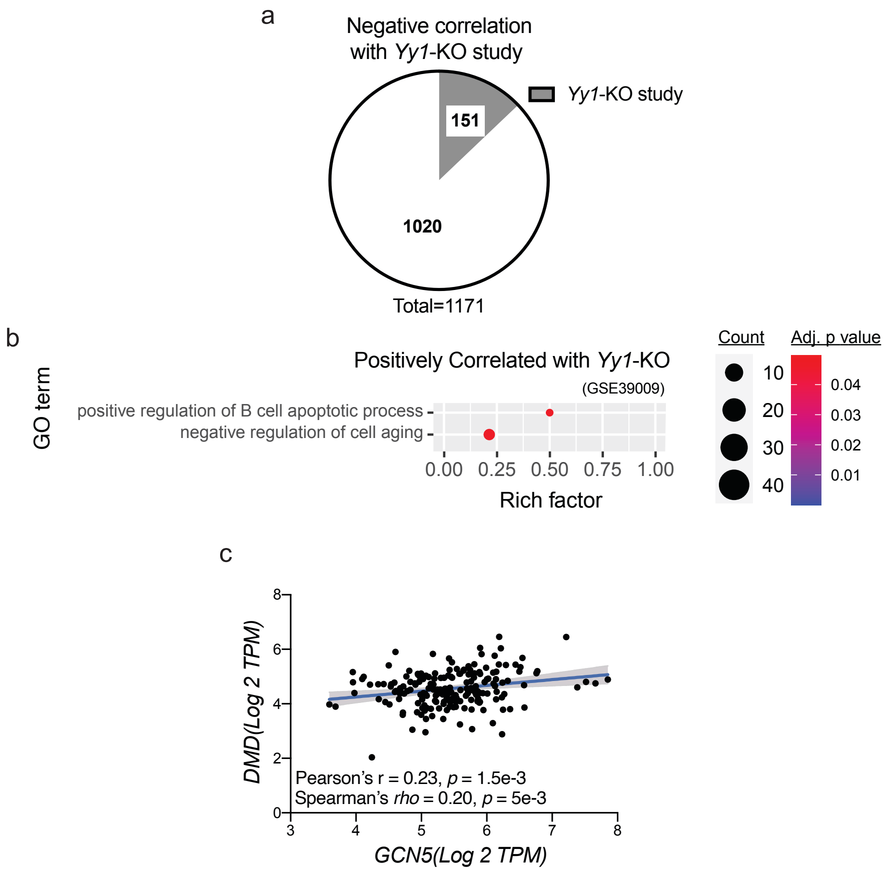
a
Negative correlation with *Yy1*-KO study
*Yy1*-KO study
151
1020
Total=1171
b
Positively Correlated with *Yy1*-KO
(GSE39009)
GO term
positive regulation of B cell apoptotic process
negative regulation of cell aging
0.00 0.25 0.50 0.75 1.00
Rich factor
Count
10
20
30
40
Adj. p value
0.04
0.03
0.02
0.01
c
DMD(Log 2 TPM)
GCN5(Log 2 TPM)
Pearson's r = 0.23, p = 1.5e-3
Spearman's rho = 0.20, p = 5e-3

**Figure S4.** YY1 regulates dystrophin and DAPC expression.

(a) Pie chart representing negatively correlated genes between *Gcn5*$^{skm-/-}$ and *Yy1*-KO (GSE39009) biosets following meta-analysis of the selected studies for downregulated and upregulated genes.

(b) Bubble plot showing the distribution and size of positively correlated over-represented gene ontology pathways between the *Gcn5*$^{skm-/-}$ differentially expressed bioset and the *Yy1-KO* study bioset (GSE39009).

(c) Correlation between *GCN5* and *DMD* gene expression as determined by RNA-Seq in human skeletal muscle from the same 192 GTEx samples (GTExPortal database v8) as in Figure 4E.

Fig S5.

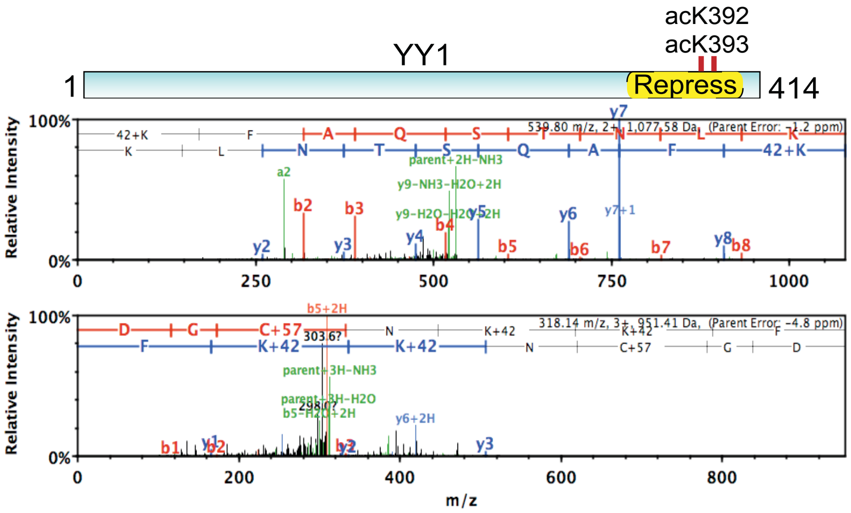

acK392
acK393
YY1
1
Repress
414
539.80 m/z, 2+, 1,077.58 Da, (Parent Error: -1.2 ppm)
318.14 m/z, 3+, 951.41 Da, (Parent Error: -4.8 ppm)
Relative Intensity
m/z

**Fig S5. Nano-LC-MS/MS analysis**

Nano-LC-MS/MS analysis showing GCN5-dependent *in vitro* acetylation of acK392 and acK393 of YY1.

#### Titles of Supplementary excel files

**Supplemental Excel File 1.** Microarray features and analysis from control and *Gcn5*$^{skm-/-}$ mice. Related to Figures 1d and 1e.

**Supplemental Excel File 2.** Differentially expressed genes and analysis from studies related to Figure 3b.

**Supplemental Excel File 3.** Differentially expressed genes and analysis from studies related to Figures 3c, S3a and S3b.

**Supplemental Excel File 4.** TRANSFAC query of factors regulating *Dmd*

**Supplemental Excel File 5.** Differentially expressed genes and analysis from studies related to Figure S4a.

#### Materials and Methods

##### Generation of Gcn5$^{skm-/-}$ mice.

For the generation of *Gcn5* floxed (*Gcn5*$^{L2/L2}$) mice that possess loxP sites flanking exons 5-6 of the Gcn5 (KAT2A) gene, genomic DNA covering the *Gcn5* locus was amplified from the 129Sv strain by using high-fidelity PCR. The resulting DNA fragments were assembled into the targeting vector designed to insert a *loxP* site upstream of exon 5, and second *loxP* site, followed by a *frt*-flanked neomycin selection cassette, downstream of exon 6 of the *Gcn5* gene.

The construct was then electroporated into 129Sv embryonic stem (ES) cells. The karyotype was verified and several correctly targeted ES cell clones were injected into blastocysts from C57BL/6J mice and resulting chimeric males had the neo cassette removed by breeding with *ACTFLPe* mice on a C57BL/6J background. Progeny were crossed to remove the *Flp*-expressing transgene and mice were subsequently bred to C57BL/6J mice for at least 10 generations. Offspring that transmitted the mutated allele, in which the selection marker was excised, and that lost the *Flp*-expressing transgene ($Gcn5^{L2/WT}$ mice) were selected, mated with human skeletal actin (HSA)-Cre mice, and then further intercrossed to generate pre-mutant $HSAcre^{Tg/0}/Gcn5^{L2/L2}$ mice. A PCR genotyping strategy was subsequently used to identify $HSAcre^{Tg/0}/Gcn5^{L2/L2}$ and $HSAcre^{0/0}/Gcn5^{L2/L2}$ mice.

**General animal phenotyping**

Animal experiments were approved by the ethics committee at the University of Ottawa with permit ID #ME-2804 and by ethics committee of the canton of Vaud, Switzerland with the Permit ID #2285. Phenotyping experiments were performed with validated Eumorphia /EMPReSS standard operating protocols (www.eumorphia.org). All mice were maintained in a temperature-controlled (23 °C) facility with a 12 hr light/dark cycle with ad libitum access to food and water. Regular chow diet (CD) was obtained from Harlan Teklad (diet 2018) contained 18.6 % protein, 44.2% carbohydrate and 6.2 % fat. Body weight (BW) was measured from 8 to 25 weeks of age. The mice were fasted 4 hrs before harvesting blood for subsequent blood measurements, and tissues for protein or RNA isolation, and histology (Champy et al., 2004, 2008).

OGTT and ipGTT were performed in overnight fasted animals. Glucose was administered by gavage at a dose of 2 g/kg BW. ipITT was done in 4h fasted animals. Insulin was injected at a dose of 0.50 U/kg of BW. Glucose measurements were performed with the ACCU-CHEK® Aviva combo system (Roche Diagnostics). Plasma insulin concentrations were measured using an ELISA for mice (Cristal Chem Inc., Downers Grove, IL).

Cold test was performed by recording body temperature using a thermometer (Bioseb) with a rectal probe (Physitemp Instruments). Mice were placed in individual cages without food and free access to water. The initial body temperature is measured before placing animals in a cold room. Mice are then placed in a cold room at 4°C. Rectal temperature is measured every hour over a period of 6 hours.

An endurance test was performed using a variable speed belt treadmill enclosed in a plexiglass chamber with a stimulus device consisting of a shock grid attached to the rear of the belt (Panlab, Barcelona, Spain). The initial velocity of the belt was 5m/min with an inclination of 5$^{o}$. The speed was gradually increased by 2m/min every 12 minutes. Exhaustion was assumed when mice received more than 5 shocks (0.1 mA) over a period of one minute. The test was considered completed after 2hrs of running. The distance traveled and time before exhaustion were recorded as maximal running distance and period.

**Downhill running experiments**

Mice were placed in a treadmill with an angle of -15 degrees and habituated to a on a variable speed belt treadmill enclosed in a plexiglass chamber with a stimulus device consisting of a shock grid (Panlab, Barcelona, Spain) for 5 minutes 1 week before the test. On the day of the test, the mice were fasted for 2hrs prior to entering the treadmill with an angle of -15$^{o}$ and were again habituated to the apparatus for 5 min. The mice then began running at a very moderate speed of 8m/min, which was then gradually increase every 5 minutes by 2m/min until the speed reached 11m/min. This speed of 11m/min was maintained for 40 min. Mice were then be rested for 5 min before a 2-minute ramp up to 11m/min for another 40 min. Mice that received 5 shocks (0.1 mA) over a period of one minute were considered exhausted and withdrawn from the treadmill.

**Eccentric Contraction of Diaphragm muscle**

Eccentric Contraction of Diaphragm muscle was performed as previously described (Addicks et al., 2018). Briefly, were triangular strips were prepared from diaphragm muscles, mounted on a custom-built force measurement apparatus and maintained under flowing physiological solution (Physiological solution 118.5 mM NaCl, 4.7 mM KCl, 2.4 mM CaCl2, 3.1 mM MgCl2, 25 mM NaHCO3, 2 mM NaH2PO4, and 5.5 mM D-glucose, bubbled with 95% $O_2$ - 5% $CO_2$. Muscle mounting was adjusted to generate maximal isometric force when stimulated by a 400 ms train of 0.3 ms square electrical pulses at 10 V and 180 Hz (701C: Electrical Stimulator; Aurora Scientific, Aurora, Canada). Force generation was measured at 5 kHz using a force transducer servomotor system (0.5N force 10mm excursion, 300C: Dual-Mode Muscle Levers; Aurora Scientific, Aurora, Canada). Diaphragm muscle was equilibrated by maintenance in physiological solution for 30 minutes with stimulated contractions every 100 seconds resulting in no decrease in contraction force over the 30-minute period. After the 30-minute equilibration muscles were subjected to eccentric contractions by stimulating as above while stretching by

10% of muscle length over a 200 ms period during the last 200 ms of each contraction. Muscle force generation was recorded during a series of 12 eccentric contractions.

**Evans blue assessment of muscle damage**

Muscle damage was assessed by injecting a 1% solution of Evans Blue dye (EBD) into the peritoneal cavity, using 1% volume to body weight, 48 hours before sacrifice. EBD was dissolved into PBS and sterilized using filters with a 0.2-µm pore size. Muscle sections from EBD-injected animals were incubated in ice-cold acetone at −20°C for 10 min, washed three times for 10 min with PBS and mounted with VECTASHIELD Mounting Medium. Microscopy images of fluorescence from EBD-positive muscle fibers were analyzed using ImageJ software.

**Creatine kinase**

Collected plasma was used for creatine kinase measurements, using the Creatine Kinase Flex Reagent Cartridge (Siemens Healthcare Diagnostics AG), along with creatinine and total protein measures on the Dimension Xpand Plus Instrument (Siemens Healthcare Diagnostics AG).

**Histology**

Muscle tissue were harvested from anaesthetized mice and immediately frozen in Tissue-TEK® OCT compound (PST) using nitrogen cooled isopentane for 2 min before being stored at -80 $^{o}$C. 8-µm cryosections fixed with 4% paraformaldehyde and stained with either haematoxylin/eosin (HE) or antibodies. For immunostainings, heat activated antigen retrieval was performed by placing sectioned tissues into a bath of pH 6.0 citrate buffer for 10 min at 65°C. Sections were then washed with PBS-0.1% tween 20 (PBST) and blocked with 10% affinipure Fab goat anti mouse IgG (Jackson Immunoresearch) in PBST for 60 min and PBST containing 2% BSA and 5% goat serum for 30min at RT. This was followed by an overnight application of mouse anti dystrophin primary antibody (Millipore, MAB1645) at 4°C followed by washing in PBST and incubation with a secondary antibody coupled Alexa-568 fluorochromes (Life technology) After washing in PBST, tissue sections were mounted using Dako mounting medium (Dako).

**Genomic profiling**

Gene regulatory networks were captured by acquiring mRNA expression patterns using microarrays (Affymetrix Mouse Gene 1.0 ST platform; https://pubmed.ncbi.nlm.nih.gov/19944634/) on 4 samples of gastrocnemius muscles taken from each experimental group. Male 6-month-old mice were fasted for 5 hours before sacrifice. Gastrocnemius skeletal muscle was immediately excised and snap-frozen in liquid nitrogen for RNA extraction. Trizol extraction of total RNA was performed for each sample. Sample concentrations were then measured by Nanodrop and cleaned up with a Qiagen RNEasy kit prior to preparation for microarray. Raw microarray data are publicly available on NCBI GEO (www.ncbi.nlm.nih.gov/gds) under the accession number GSE158883.

**Correlation Engine**

In order to validate our results and find associations by matching and integrating our transcriptomic data with publicly available genomic knowledge, we used Illumina BaseSpace™ Correlation Engine (BSCE, Illumina, San Diego, CA, https://sapac.illumina.com/products/by-type/informatics-products/basespace-correlation-engine.html). The *Gcn5*$^{skm-/-}$ *vs* wildtype differentially expressed geneset was used as the input bioset and correlated to the 142,219 ontologically tagged biosets comprising 23,529 Curated Studies on 29 June 2020 (Kupershmidt et al., 2010).

Bioset data stored in the BSCE undergo several preprocessing, quality control, and organization stages. Quality examinations ensure the integrity of the samples and datasets and include evaluations of pre- and post-normalization boxplots, missing value counts, and p-value histograms after statistical testing with a false discovery rate analysis to establish whether the number of significantly altered genes is larger than the number anticipated by chance.

The rank-based, nonparametric Running Fisher algorithm implemented within BSCE was used to compare the transcriptomic bioset derived from *Gcn5*$^{skm-/-}$ *vs* wildtype mice to other transcriptomic biosets in the database. The Running Fisher algorithm computes statistical significance of similarity between ranked fold change values of two bioset gene lists using a Fisher's exact test. This normalized ranking approach enables comparability across data from different studies, platforms, and analysis methods.

Based on the bioset-bioset correlations of disease *vs* normal experimental designs (31,955 biosets) and associated phenotype tags, the BSCE Disease Altas was used to identify highly correlated musculoskeletal diseases. The BSCE Knockout Atlas was used to identify highly correlated transcriptomic profiles for available gene knockout models. The BSCE Meta-Analysis application was used to aggregate, score and rank genes across the *Gcn5*$^{skm-/-}$ *vs* wildtype bioset and select highly ranked musculoskeletal disease biosets in BSCE Curated Studies to assess common and differential RNA expression activities. Over-representation gene ontology pathway analysis was performed for the *Gcn5*$^{skm-/-}$ differentially expressed geneset and the genesets obtained following meta-analysis (InnateDB) (Breuer et al., 2013).

**Human Minimum Feret Diameter and Gene Expression Correlation**

The correlation analysis in 119 human skeletal muscles were performed using RNA-seq dataset belonging to GSE111017 (NCBI GEO, www.ncbi.nlm.nih.gov/gds) (Migliavacca et al., 2019). All downloaded data sets were normalized using TMM normalization method. Finally, all data were converted to Z-score to avoid batch effects. Pathway enrichment plots and scatter plots were plotted using the ggplot2 R package as the Z-score of TMM-normalized TPM.

Images of human cadaver muscle sections and RNA-seq data for corresponding muscle tissue were obtained from publicly available datasets from the human Genotype-Tissue Expression (GTEx) consortium (GTExPortal database v8). Fiber size quantification was performed in ImageJ for 192 images (~25% of all images). A total of 250 fibers were measured for minimum feret diameter per image. The mean minimum feret diameter for each image was correlated with corresponding *YY1* mRNA expression (Log2 TPM). GTEx expression data for *YY1* is available here: http://www.gtexportal.org/home/gene/YY1. GTEx histology images and pathological notes are available using the GTEx Histology Image Viewer: https://gtexportal.org/home/histologyPage.

**EMSA**

YY1 and YY1 with lysines 392 and 393 mutated to arginine or glutamine were expressed from plasmid DNA using the TnT® Quick Coupled Transcription/Translation System (Promega, L1170) according to manufacturer's protocol. Oligonucleotides containing YY1 binding sites (CGCTCCCCGGCCATCTTGGCGGCTGGTGG, CCACCAGCCGCCAAGATGGCCGGGGAGCG) or mutated sites (CGCTCCGCGATTATCTTGGCGGCTGGTGG, CCACCAGCCGCCAAGATAATCGCGGAGCG) were suspended in water and annealed by heating to 95˚C then cooling to 21˚C at a rate of 0.5 ˚C per minute in a thermocycler. Radiolabeled oligonucleotide dimers were prepared by incubating annealed oligonucleotides with 32P-ATP in the presence of T4 PNK (New England Biolabs, M0201) according to manufactures protocol and purified using Micro Bio-Spin P-30 Tris Chromatography Columns (Bio-Rad, 7326250). YY1 was bound to oligonucleotide dimers in a solution of 10 mM Tris pH 7.5, 60 mM KCl, 2 mM MgCl2, 0.5 mM DTT, 10% glycerol, 0.2 mg/ml BSA and ½ vol/vol of reticulate lysate reaction in a 15 ul reaction. For each reaction 50 fmol of labeled probe was added. 5 pmol of unlabeled dimers were added to specified reactions. All reactions also contained 0.1 ug of sonicated salmon sperm DNA (ThermoFisher, 15632011) to reduce non-specific binding. Binding reactions were incubated at 25 ˚C for 30 minutes and loaded directly on a 6% acrylamide non-denaturing tris borate gel in the absence of EDTA. Gels were run at 60 V for 1 hour, backed with Whatman paper, and dried for 2 hours at (80 ˚C) in a gel dryer under vacuum. Dried gels were exposed to film at -80 ˚C overnight with an intensifying screen.

**YY1 overexpression in myoblasts**

YY1 retrovirus was prepared by transfecting 293T cells with plasmids containing YY1 under control of the EF1a promotor (parent plasmid was pLV-EF1a-IRES-Neo, a gift from Tobias Meyer (Addgene plasmid # 85139 ; http://n2t.net/addgene:85139 ; RRID:Addgene_85139), and helper plasmids pMDL, pREV, and pVSVG using the calcium chloride method. Myoblasts were infected with a m.o.i. of ~0.001 by adding viral supernatant at a 1/1 ratio to growth media and incubated overnight in a 35 mm dish. Myoblasts were split to 2 x 10 cm dishes and after a further 24 hours selected for neomycin expression by addition of G418. After 4 to 5 days isolated individual colonies were transferred to wells in a 48 well dish, expanded and assessed for YY1 expression. The four highest YY1 and neomycin expressing colonies (and neomycin expressing controls), as determined by qPCR, were plated into 35 mm wells and grown to 33 % confluence, transferred to differentiation medium (DMEM with 5% horse serum) for 7 days and collected for qPCR and western blot.

**RNA isolation and qPCR**

Cells were collected by trypsinization and centrifugation, then washed once with 10% BSA in PBS and once with PBS. RNA was prepared using EZ-10 Spin Column Total RNA Miniprep Kit (BioBasic, BS1361(SK8655)) according to manufacturer's instructions. RNA was quantified using NanoDrop (Thermo) and for each sample 1 ug RNA was converted to cDNA using ProtoScript II Reverse Transcriptase (New England Biolabs M0368) in a 20 ul reaction. 0.5 ul of RT was used per 10ul qPCR reaction (Luna® Universal qPCR Master Mix).

Total RNA from mouse muscle was extracted using TRIzol and then transcribed to complementary DNA using the QuantiTect Reverse Transcription Kit (Qiagen). Expression of selected genes was analyzed using the LightCycler 480 System (Roche) and SYBR Green chemistry. All quantitative polymerase chain reaction (PCR) results were presented relative to the mean of *36b4*, *B2m*, and/or *Gapdh* ($\Delta\Delta C_t$ method).

**Primer sets for qRT-PCR analyses:**

| Gene | Description | Forward Primer | Reverse Primer |
|---|---|---|---|
| *36b4* | Ribosomal protein, large, P0 | AGATTCGGGATATGCTGTTGG | AAAGCCTGGAAGAAGGAGGTC |
| *B2m* | Beta-2 microglobulin | TTCTGGTGCTTGTCTCACTG | TATGTTCGGCTTCCCATTCT |
| *Gapdh* | Glyceraldehyde-3- phosphate dehydrogenase | TGTGTCCGTCGTGGATCTGA | CCTGCTTCACCACCTTCTTGAT |
| *Dmd* | dystrophin exon 1 (dmd) 427muscle isoform | TCTCATCGTACCTAAGCCTC | CAGTGCCTTGTTGACATTGTTCAG |
| *Utrn* | Utrophin | GCCCTCCCTGCAGATTATTTG G | CTGTCCAGTTGACCTTTGATACTCTTC |
| *eMHC* | embryonic myosin heavy chain | TGAAGAAGGAGCAGGACAC | CACTTGGAGTTTATCCACCAGATCC |
| *Maged1* | MAGE Family Member D1 | CAAGAGGACCCGCAAGGTT | GCCTTTGATCCCCACTGTTG |
| YY1 | YY1 Transcription Factor | GAAGCAGGTGCAGATCAAGACCC | GAGAGGTCAATGCCAGGTATCCC |
| Gcn5 | Lysine Acetyltransferase 2A (General Control Of Amino Acid Synthesis Protein 5-Like 2) | GGAAGGCGCAAGTCCGGG | GCTGGAGGTCCATGCGGG |
| Sgcb | Beta-Sarcoglycan | ATCCCCATCGATGAGGACCGG | CCCATTTGGCCCAATGCGG |
| Sgcd | Sarcoglycan Delta | CCACAGGAGCACCATGCCC | CTGAGTCTCCTTCTAGCTTCAGACCC |
| Sntb1 | Syntrophin Beta 1 | ACGGAGCAGACCTGCGGG | CATCCAATCTCAGACACCGGG |
| Ttn | Titin | GCCAACAGTGGACGATACTCCC | CTCTGGATTTCGGCTCCATCCC |
| Actn1 | Actinin Alpha 1 | CCTGGATCCGGCCTGGG | TCTCTGGCTTGGCCAAGCG |

| | | | |
|---|---|---|---|
| Actn2 | Actinin Alpha 2 | CTCGACCCGGCCTGGG | TTTCCCCGGTCAGGTTTGGG |
| Lama2 | Laminin Subunit Alpha 2 | CACATGGTGGCAGAGTCCC | CCAAAATCCAGTTTCCAGGCC |
| Lamc1 | Laminin Subunit Gamma 1 | CCAGACTATGCTGGCCGGG | CTGGTGTGGAACTTGAGGCG |
| Lamc2 | Laminin Subunit Gamma 2 | CAGCCTCAGTACCACGCCC | CTGCTGTGCCTTCCTTTTCCC |
| Itga4 | Integrin Subunit Alpha 4 | AGGGATAACCAGTGGCTGGG | GACGTAGCAAATGCCAGTGGG |
| Ckm | Creatine Kinase, M-Type | GCCATGGCGGCTACAAACCC | GCGGAGGCAGAGTGTAACCC |
| Pcaf | P300/CBP-associated factor | CGAAGCTGTAGCCATGCCC | CTCAAATGGCGGCTTCTTCTCC |

**Western blotting**

Cells were collected by trypsinization and centrifugation, washed once with 10% BSA in PBS and once with PBS. Cells were lysed in cold 0.5% Triton Buffer (0.5% tryton-X, 5 mM Tris, pH7.6, and 150 mM NaCl), quantified using the Bradford method and diluted to 0.1 mg/ml with Triton Buffer. 5ug of protein was electrophoresed on a 6% SDS PAGE gel with 0.05% TCE and UV stain-free imaged. Proteins were blotted to PVDF membranes blocked with milk and detected using mouse anti-dystrophin (Millipore, MAB1645), anti-HSP90 (BD Biosciences), anti-YY1 (Santa Cruz, H-10), anti-acetylated-Lysine (Cell Signaling #9441), anti-GCN5 (Santa Cruz, sc-20698), anti-GST (Abcam, ab9085) and anti-mouse HRP secondary antibody (NEB, 7076). Antibody detection reactions were developed by enhanced chemiluminescence (Advansta) and imaged using the ChemiDocTM Touch Imager system (Bio-Rad Laboratories Ltd. – Catalog No.1708370).

Muscle tissue was snap frozen using liquid nitrogen and pulverized at -80°C. Powdered muscle was then homogenize using Qiagen Tissue Lyser for 3 min at 50 Hz in RIPA buffer (20 uL/mg of tissue) with protease inhibitor cocktail (Roche, 11836153001). Samples were quantified using the Bradford method. 20ug of protein was electrophoresed on 5-8% gradient gel. Proteins were blotted to PVDF membranes blocked with BSA and detected using mouse anti-dystrophin (Millipore, MAB1645), anti-HSP90 (BD Biosciences), anti-acetylated-Lysine (Cell Signaling #9441) primary antibodies and anti-mouse HRP secondary antibody (NEB, 7076).

**Cell Culture and treatments of C2C12 myoblasts**

C2C12 mouse muscle derived myoblasts (CRL-1772TM, ATCC) were cultured in Dulbecco's modified Eagle's medium (DMEM) including 4.5 g/L glucose, 20% fetal calf serum and 50 µg/ml gentamicin. Differentiation of C2C12 cells into myotubes was induced over four days in Dulbecco's modified Eagle's medium (DMEM) including 4.5 g/L glucose, 2% horse serum and 50 µg/ml gentamicin. Cells were tested for mycoplasma using Mycoprobe (#CUL001B, R&D systems). *Gcn5* knockdown was performed using either adenovirus infection (Addgene) or cell transformations with *Gcn5* shRNA (Santa Cruz) using jetPEI DNA transfection kit (Polyplus), according to manufacturer's instructions.

**Plasmids and Recombinant Adenoviruses for in vitro acetylation assays**

A plasmid expressing GCN5::FLAG was obtained from Addgene (#14106). To generate the plasmid expressing GST::YY1 (pGEX-5X-1-GST-YY1), the CDS of 3 YY1 fragments (F1: 1-190; F2:

180-300; F3: 300-414) were amplified from murine muscle cDNAs and ligated to pGEX-5X-1 plasmids.

**In Vitro Acetylation Assays**

In vitro acetylation and deacetylation assays were performed as originally described (Rothgiesser et al., 2010). In brief, 1 µg of recombinant YY1 protein obtained from BL21 strain was incubated with 500 ng of recombinant GCN5 in acetylation buffer (150 µM acetyl-CoA, 50 mM Tris-HCl [pH 8], 100 mM NaCl, 10% glycerol, 1 mM phenylmethylsulfonyl fluoride [PMSF], 1 mM dithiothreitol [DTT], 1 µg/ml bepstatin, 1 µg/ml leupeptin, 1 µg/ml pepstatin, and 1 mM sodium butyrate) for 1 hr at 30°C. After incubation, samples were resolved with SDS-PAGE sample loading buffer and analyzed by western blot or with nano liquid chromatography tandem mass spectrometry (LC-MS/MS) to define the acetylated residues.

**Mass Spectrometry**

Gel lanes were cut into pieces followed by an in-gel digestion with endoproteinase Glu-C or trypsin. These peptide digests were then resuspended and analyzed using nano LC-MS/MS with an Orbitrap Elite Mass Spectrometer (Thermo Fischer Scientific) coupled to an ultraperformance LC system (Thermo Fischer Scientific Ultimate 3000 RSLC). Data analysis was performed with Proteome Discoverer (v. 1.3), and searches were performed with Mascot and Sequest against a mouse database (UniProt). Data were further processed, inspected, and visualized with Scaffold 4 (Proteome Software, Inc, USA).

**Statistics**

Differences between two groups were assessed using two-tailed t tests. Differences between more than two groups were assessed using one-way analysis of variance (ANOVA). To compare the interaction between two factors, two-way ANOVA tests were performed. ANOVA, assessed by Bonferroni's multiple-comparison test, was used when comparing more than two groups. GraphPad Prism 5 was used for all statistical analyses. All P values <0.05 were considered significant. *$P \leq 0.05$, **$P \leq 0.01$, and ***$P \leq 0.001$.
